# A lightweight data-driven spiking neural network model of *Drosophila* olfactory nervous system with dedicated hardware support

**DOI:** 10.1101/2023.10.12.560618

**Authors:** Takuya Nanami, Daichi Yamada, Makoto Someya, Toshihide Hige, Hokto Kazama, Takashi Kohno

## Abstract

Data-driven spiking neural network (SNN) models are vital for understanding the brain’s information processing at the cellular and synaptic level. While extensive research has focused on developing data-driven SNN models for mammalian brains, their complexity poses challenges in achieving precision. Network topology often relies on statistical inference, and the functions of specific brain regions and supporting neuronal activities remain unclear. Additionally, these models demand significant computational resources. Here, we propose a lightweight data-driven SNN model that strikes a balance between simplicity and reproducibility. We target the *Drosophila* olfactory nervous system, extracting its network topology from connectome data. The model implemented on an entry-level field-programmable gate array successfully reproduced the functions and characteristic spiking activities of different neuron types. Our approach thus provides a foundation for constructing lightweight *in silico* models that are critical for investigating the brain’s information processing mechanisms at the cellular and synaptic level through an analysis-by-construction approach and applicable to edge artificial intelligence (AI) systems.

## 1 Introduction

Elucidating the mechanisms underlying information processing in the brain represents a great challenge in neuroscience. In parallel to collecting data with experiments, building brain models has proven to be a powerful approach to enable *in silico* analysis and provide a framework for understanding information processing in the brain. Macroscopic models [1][2][3][4] describe information flow at the functional level and present an overview of neural processing. In contrast, spiking neural network (SNN) models emulate the brain at the cellular and synaptic level and provide their *in silico* counterparts, which are more tractable and easier to manipulate. From an engineering perspective, properly built SNN models are expected to be capable of intelligent information processing equivalent to the brain. Silicon neuronal network (SiNN) chips, which are highly power-efficient neuromorphic hardware optimized for SNN models, have already been developed [5][6][7]. Therefore, they have great potential for next-generation artificial intelligence (AI) applications.

The structure of the brain is highly diverse, which makes it demanding to capture the comprehensible rules about the network topology. In addition, a wide variety of neuronal and synaptic properties has been reported. The data-driven approach intends to replicate the brain by semi-automatically incorporating vast amounts of anatomical and physiological data. Several large-scale data-driven SNN models [8][9][10] that reproduce a part of the mammalian cortex and hippocampus have been built. They were designed to replicate the network topology, neuronal anatomy and electrophysiology, and synaptic properties, and they successfully reproduced the characteristic spiking activities seen in the target regions. However, in mammalian brains, the considerable number of neurons makes it challenging to measure the exact connection topology between the neurons. Hence, the network topology was inferred based on statistical data. In addition, because each brain region closely interacts with various other brain regions, it is not trivial to understand the specific function of the target region. Generally, data-driven models employ the ionic-conductance-based neuronal models, which can reproduce arbitrary electrophysiological properties but incur enormous computational costs. For example, the model in [9] runs on a supercomputer consisting of 3488 processors, and its simulation speed is 1600 times slower than real-time. Moreover, these models are not suitable for implementation on SiNNs because they involve complex calculation processes that require enormous circuit resources.

In this study, we built a data-driven SNN model for the olfactory nervous system of *Drosophila melanogaster* (fruit fly). The system is a relatively small (∼2200 neurons) network having a known function, whose complete network topology, or connectome, is available. The electrophysiological activity of neurons was reproduced by using the piecewise quadratic neuron (PQN) model, which is a lightweight neuron model suitable for digital arithmetic circuit implementations [11][12][13][14][15][16].

The PQN model was adopted to reduce the computational cost and enable the SNN model to be run on a SiNN chip. It focuses on reproducing the key dynamics behind neuronal activities with lightweight calculations. The model is designed using the dimension reduction techniques of nonlinear dynamics such that the dynamical structure of the activity of the target neuron is preserved. Unlike integrate-and-fire (I&F) based models, such as the leaky I&F model, Izhikevich (IZH) model [17], and adaptive exponential I&F model [18], the dynamics in the neuronal spike are not replaced by a resetting of the membrane potential. I&F-based models are generally more lightweight than the PQN model. However, they have been reported to have limitations in the reproducibility of neuronal activities. For example, because their spike amplitudes are fixed, they cannot reproduce the propagation of spike intensity observed in some brain regions including the hippocampus [19]. In addition, the IZH model can only reproduce spiking within a limited range of stimulus intensities [11]. Furthermore, the phase-resetting curve of the Class II mode in Hodgkin’s classification [20] of the IZH and AdEx models differs from the typical shape [16]. In addition to the aforementioned advantages, the PQN model supports efficient implementation on digital arithmetic circuits. Thus, the SNN model can be executed efficiently (power and speed) with a SiNN on field-programmable gate arrays (FPGAs) and application-specific integrated circuits (ASICs). The results in this study were obtained using a SiNN on an entry-level low-cost FPGA chip to demonstrate its potential for low-power brain-morphic AI applications.

The fruit fly brain comprises 100,000 neurons. Moreover, its connectome was recently revealed [21]. It is compact compared to the mammalian brain but capable of complex information processing. Its olfactory nervous system consists of brain regions including the antennal lobe and the mushroom body, the anatomy and physiology of which have been widely studied [22][23]. The function and activity of each type of neuron in these regions are better characterized in the context of sensory input and behavioral output than those of the mammalian cortex and hippocampus, enabling us to adequately verify the reproducibility of the model. However, previous modeling studies [24][25][26] (not data-driven) of the olfactory nervous system used simplified I&F-based neuron models, which did not fully reproduce the electrophysiological properties of each type of neurons. Specifically, they did not reproduce the characteristic spiking activities seen in the olfactory nervous system including (1) odor-evoked oscillatory firing in the projection neurons (PNs) and local neurons (LNs) [27], (2) absence of oscillations in Kenyon cells (KCs) [28], (3) different contributions of LN subclasses to the formation of oscillations [27], and (4) temporal dynamics of firing in mushroom body output neurons (MBONs) [29]. Thus, it is uncertain whether they accurately capture information processing mechanisms in the olfactory nervous system. More sophisticated, ionic-conductance-based SNN models of the insect brain [30][31] had been built for the antennal lobe of locust. However, they were not data-driven and did not reproduce most of the aforementioned characteristics of spiking activities. This is likely because they modeled only PNs and LNs, and also lacked electrophysiological data on identified neurons. Here we built a model of a fly olfactory system incorporating the connectome data as well as neuronal and synaptic electrophysiological properties of neurons. Our model successfully reproduced not only the aforementioned characteristic spiking activities (1)–(4) of the constituent cells, but also olfactory associative plasticity, the primary function of the olfactory system. Although we did not intend to implement every single known neuron or connection in our model, this study lays a foundation for building lightweight data-driven SNN models and is expected to aid in understanding the brain and developing brain-morphic AI systems.

## 2 Results

### 2.1 Overview of the model

This section provides an overview of the proposed network model. The details of the model and its background, including the justification of the choice of neurons, are explained in the Methods section. The model comprises an antenna, the antennal lobe, and the mushroom body (Fig. 1). The antenna contains olfactory receptor neurons (ORNs), and the antennal lobe contains LNs and PNs [34]. KCs, an anterior paired lateral neuron (APL), and MBONs are present in the mushroom body [35]. The two MBONs, MBON-*α*1 and MBON-*α*3, project to SMP354 neuron, whose excitatory activity can trigger a series of olfactory approach behaviors including upwind steering and locomotion [36]. As mentioned in Supplementary Note 1, whereas ORNs, PNs, KCs, and MBON-*α*3 are excitatory, LNs, APL, and MBON-*α*1 are inhibitory in nature. The numbers in Figure 1 represent the numbers of neurons implemented in the model. Synaptic connections were determined based on the connectome database hemibrain v1.2.1 [21][37], which comprehensively describes the connections between neurons in the brain of adult *Drosophila melanogaster*. However, owing to the lack of information about the types of LNs in the connectome database, the connections of LNs were determined based on a different study [32] (see Methods section).

**Figure 1:**
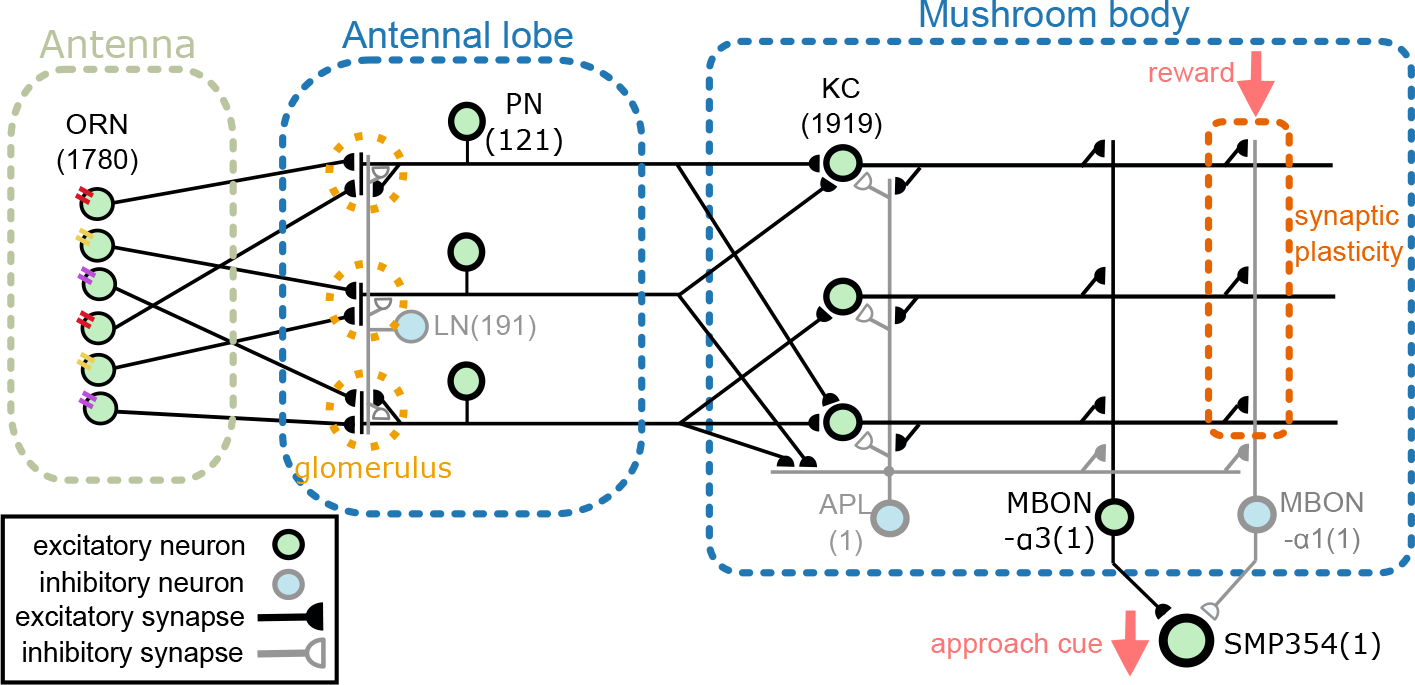
Network overview. The network comprises an antenna, the antennal lobe, and the mushroom body. ORNs, PNs, KCs, and MBON-*α*3 are excitatory neurons, whereas LNs, APL, and MBON-*α*1 are inhibitory neurons. SMP354 neuron receives excitatory and inhibitory input from the two MBONs and produces the approach cue. KC*>*MBON-*α*1 synapses can express synaptic plasticity, which is driven by the reward signal.

Odors are first detected by ORNs on the antenna. ORNs express one of the olfactory receptors (ORs), each possessing different odor selectivity. ORNs extend their axons to the antennal lobe and project to PNs and LNs. As the firing activities of ORNs are used as the input data, ORNs are not modeled. LNs mediate lateral interaction within the antennal lobe, and PNs send output to KCs and APL in the mushroom body. KCs provide convergent projections to MBON-*α*1 and MBON-*α*3, and APL delivers inhibitory feedback to neurons in the entire mushroom body. When odors are given, ORNs, PNs, and KCs fire in turn, and the populations of the firing KCs encode individual odors. Before learning, both excitatory and inhibitory MBONs fire when given an odor; hence, their outputs cancel with each other, and SMP354 neuron does not fire. When a reward stimulus (upper right in the figure) is given, the intensities of KC*>*MBON-*α*1 synapses (the synapses from KC to MBON-*α*1) that had been firing within five seconds are weakened from 1 to 0.25. This learning by long-term depression (LTD) leads to the selective inactivation of MBON-*α*1 to the odor preceding the reward stimulus, causing selective activation of SMP354 neuron to that odor. As SMP354 neuron triggers approach behaviors, the fly approaches the odor associated with the reward stimulus.

We prepared electrophysiological data of each neuron and tuned the PQN models to replicate them. For LNs and KC, data recorded in previous studies [32][33] were used. The detailed procedures for the data acquisition from PNs and MBONs are described in the Methods section. Owing to the lack of data on APL and SMP354 neurons, only the modeling results are shown. Since we did not have data on MBON-*α*3, we used the one on MBON-*α*1. The PQN model was used to model the neurons, the details of which are shown in the Methods section. Figure 2 illustrates the responses of the somatic membrane potentials *in vivo* (red) and those of the PQN models on FPGA (blue). The black plots are the step input currents, whose unit in the recording is pA. The FPGA simulation results have no physical unit. Although a variety of LN subclasses were observed [38][32], we employed four electrophysiologically identified subclasses reported in [32]. They are Krasavietz class1, Krasavietz class2, NP1227 class1, and NP2426 class1; we fitted PQN models to each of them. The parameters of the PQN model were automatically determined using a fitting method [14][15] based on the differential evolution algorithm [39]. Detailed activities of each neuron are shown in Supplementary Figures 1-7.

**Figure 2:**
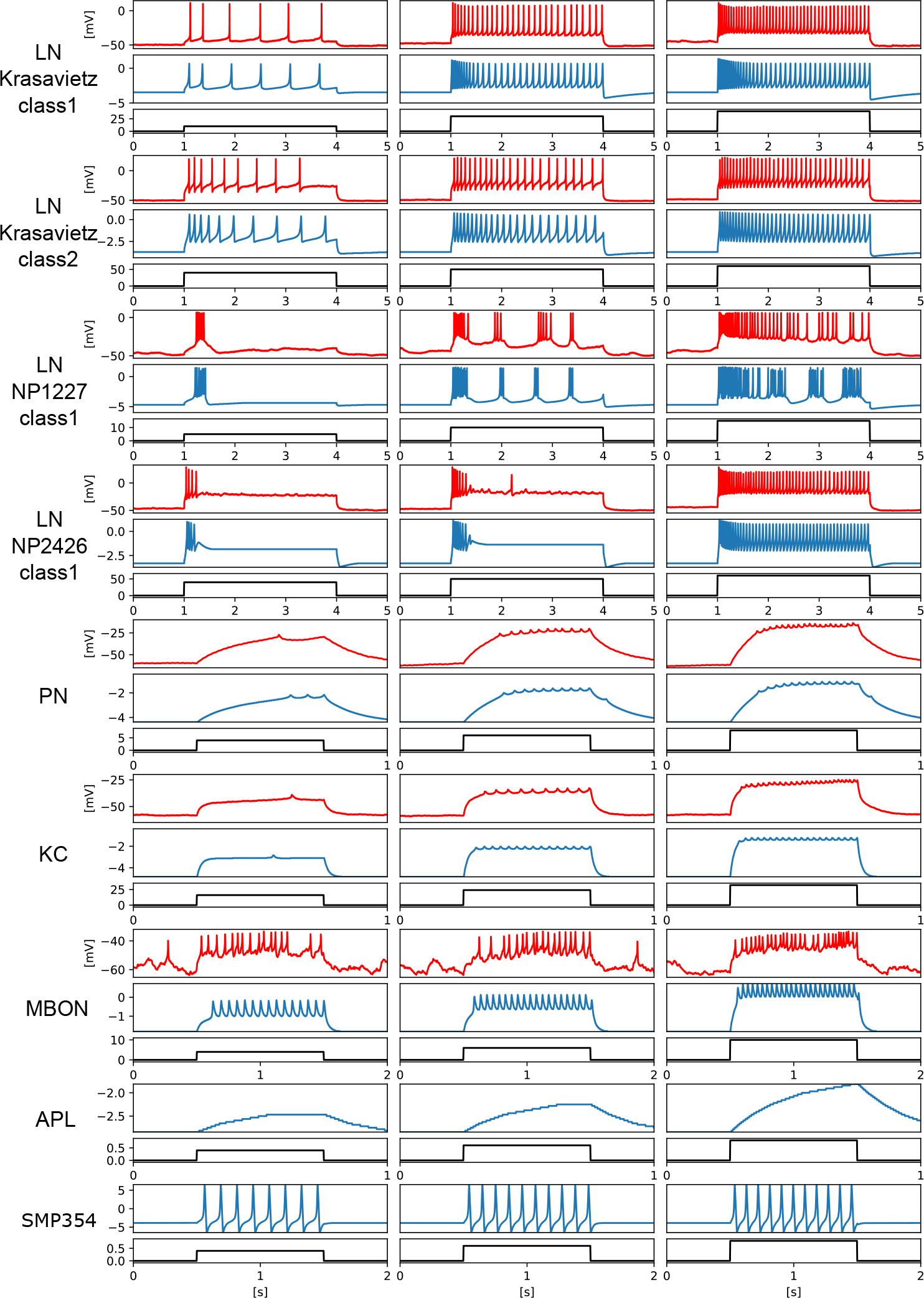
Electrophysiological properties of somatic membrane potentials of the *in vivo* data (red) and the simulated results of the PQN models *in silico* (blue) in response to step stimulus inputs (black). We conducted recordings from PNs and MBONs in this study. The data of the KC and four subclasses of the LNs are from previous studies [32][33]. As there is no recorded data of APL and SMP354 neuron, we only show the simulation results.

### 2.2 Olfactory associative learning

Flies are capable of olfactory associative learning where they remember the odor associated with the reward. One of the main goals of our model is to reproduce the neuronal mechanisms underlying this learning. Figure 3a shows a portion of the raster plots for ORNs, PNs, and KCs. Every five seconds, one of the six odorants, 3-octanol, cis-3-hexenol, cyclohexanone, 2,3-butanedione, 2-hexanol, and ethyl butyrate, was applied in turn for one second. As the responses propagate from ORNs to PNs to KCs, a smaller number of neurons are activated. These have also been observed in the olfactory nervous systems in multiple species [40][28]. This sparse activity of KCs suggests that individual odors are represented by a small number of KCs, which in turn allows flies to selectively identify the odor associated with the reward.

**Figure 3:**
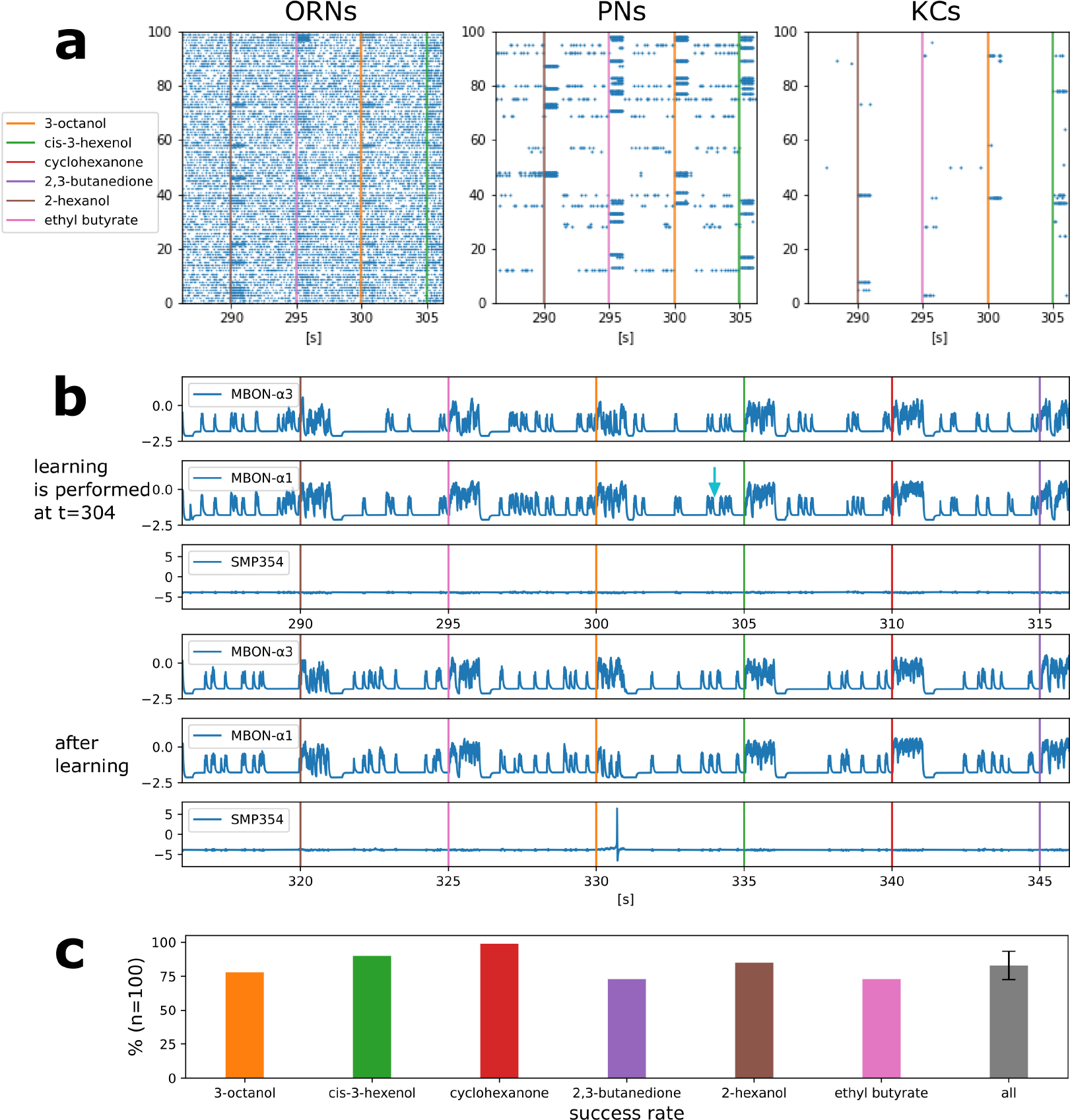
Activities of each type of neuron and the success rate of olfactory associative learning. **a** Parts of the raster plot of ORNs, PNs, and KCs. The horizontal axis represents time and the vertical axes represents indices of the neurons, respectively. The colored bars represent the onset of the one-second odorant applications. The blue dots represent spikes. **b** Waveforms of somatic membrane potentials of MBON-*α*3, MBON-*α*1 and SMP354 neurons during and after learning. The light blue arrow indicates the timing of the reward signal that triggered learning. **c** Success rates of olfactory associative learning for six odorants. Error bars represent standard deviation over the success rates of the six odorants.

Figure 3b shows the activities of MBONs and SMP354 neuron before and after olfactory associative learning *in silico* (FPGA). The application of 3-octanol was followed by a reward signal at t = 304. This resulted in LTD at KC*>*MBON-*α*1 synapses, the presynaptic neurons of which fired in the previous five seconds. Subsequently, MBON-*α*1 became selectively unresponsive to 3-octanol, whereas MBON-*α*3 remained responsive to all odorants. Consequently, SMP354 neuron that receives excitatory input from MBON-*α*3 and inhibitory input from MBON-*α*1 fires only when 3-octanol is applied.

Figure 3c shows the success rates of olfactory associative learning for individual odors. Each set of experiments comprised one associative learning and ten trials. In each trial, all six odors were applied sequentially in a unique order. A trial was considered successful when SMP354 neuron responded solely to the learned odor. Ten sets of experiments comprising 100 trials were conducted for each odor, and the probabilities of success were calculated. This model achieved an average success rate of 83.0%. The variation in the results of each trial originates from the variable input spike streams from ORNs. The activities of ORNs were calculated based on the Poisson process (details are provided in the Method section).

### 2.3 Oscillations in the antennal lobe

In order to test whether our model is applicable to known activity dynamics observed in the *Drosophila* olfactory system other than plastic changes induced by learning, we next focused on neuronal oscillations. Neuronal oscillations are widely observed in the olfactory nervous system of insects and are believed to be important in odor information processing [41][42]. Oscillations have also been reported [27] in the PNs of *Drosophila*, which are absent without odors or when LNs are inactivated. A similar oscillatory behavior was observed in our model. Whereas [27] measured the local field potential (LFP) caused by the synaptic currents of PNs, we calculated a virtual LFP by averaging the synaptic currents for each type of neuron. Figure 4a shows the virtual LFP of PNs, LNs, and KCs when 3-octanol was applied, where clear oscillations can be seen in PNs and LNs. The peak amplitudes of their frequency spectra were estimated to clarify their oscillatory nature. Figure 4b shows the power spectra (Fourier transform), which have the peak at approximately 20–30 Hz. Here, the odor was applied for ten seconds. Following this, we applied 3-octanol twenty times and plotted the averaged values of the peak power on a logarithmic scale (Fig. 4c). The peak power of LNs was considerably higher than that of PNs; whereas, the peak power of KCs was much smaller than that of PNs, which is consistent with [28] reporting no clear oscillations in the membrane potentials of the *Drosophila* KCs. In addition, the peak power when odor was not given was much lower than that under normal conditions, which is consistent with the results in [27].

**Figure 4:**
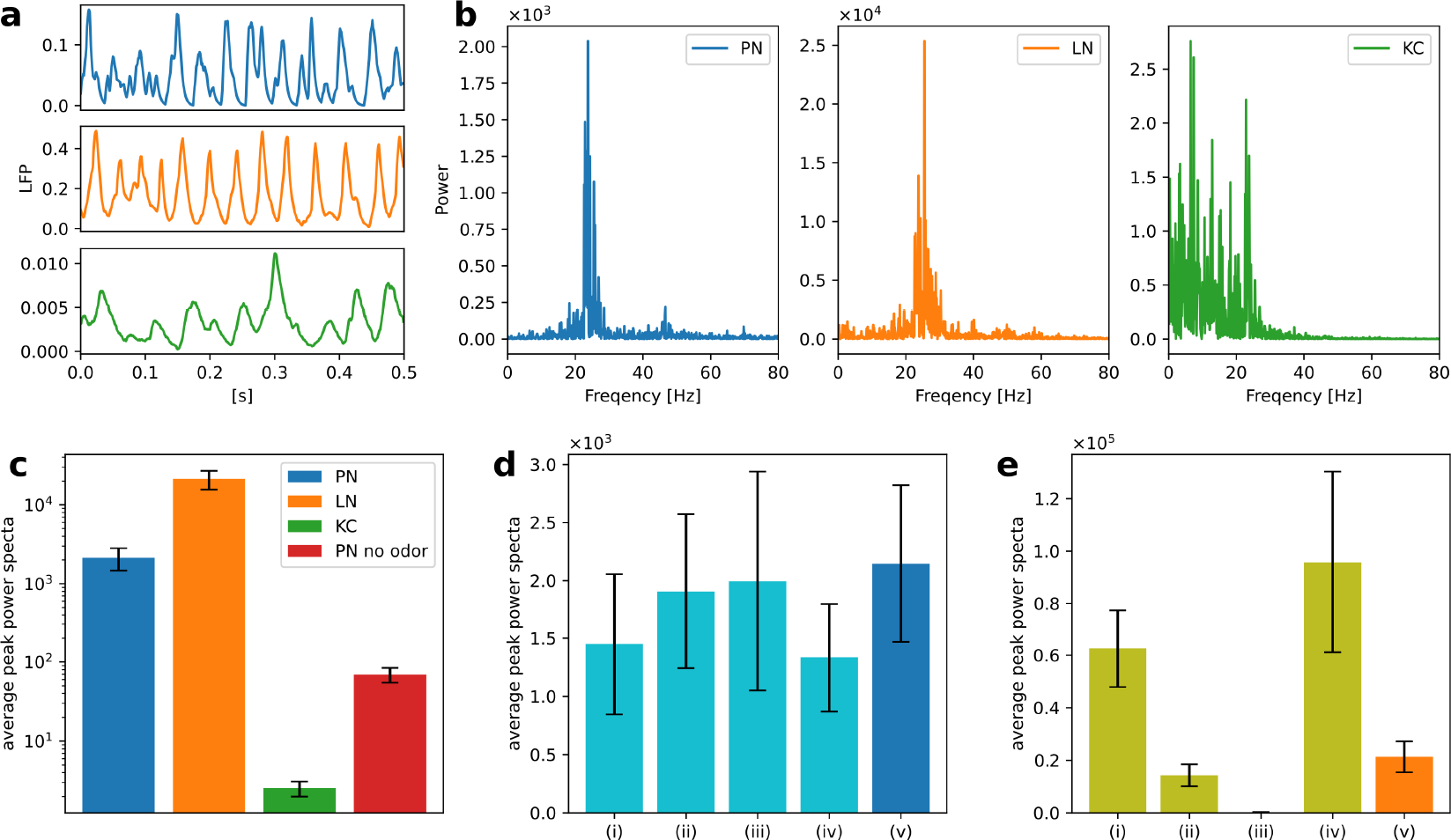
Neuronal oscillations in PNs, LNs, and KCs. **a** Examples of the virtual LFP for each type of neuron. **b** Power spectra of the virtual LFPs of PNs, LNs, and KCs. 3-octanol was applied for ten seconds. **c** Averages of the peak power spectra were plotted on a logarithmic scale graph. Average of the peak power spectra of PNs in the absence of the odor was also plotted. Error bars represent standard deviation over 20 trials. **d** Averages of the peak power spectra when one of the four types of LNs was inactivated. LNs of Krasavietz class1 (i), Krasavietz class2 (ii), NP1227 class1 (iii), and NP2426 class1 (iv) were inactivated, respectively. The results obtained without inactivation are also plotted for comparison (v). Error bars represent standard deviation over 20 trials. **e** Averages of the peak power spectra of each LN subclasses. Krasavietz class1 (i), Krasavietz class2 (ii), NP1227 class1 (iii), and NP2426 class1 (iv). The average peak power spectra of the virtual LFPs over all LNs were also plotted for comparison (v). Error bars represent standard deviation over 20 trials.

The previous study [27] selectively inactivated the synaptic output of NP1227 class1 and NP2426 class1 LNs in turn, and reported that the oscillations of PNs were attenuated only when NP2426 class1 was inactivated. We inactivated each subclass of LNs in turn and plotted the average of their peak spectra of the oscillations of PNs (Fig. 4d). Here, inactivation of LNs was performed by forcing the stimulus input to the LNs to zero. 3-octanol was applied five times for each condition. The peak power was significantly attenuated when NP2426 class1 but not NP1227 class1 was inactivated. This is consistent with the experimental results in [27]. The second and third largest attenuation was observed following the inactivation of Krasavietz class1 and class2, respectively.

We also calculated the virtual LFP for each LN subclass under the normal condition. Fig. 4e shows the average peak power when 3-octanol was applied twenty times. The peak power of NP2426 class1 was the largest, indicating that it was the primary source of oscillation in LNs. The peak powers of Krasavietz class1 and Krasavietz class2 are the second and third largest, respectively. There are almost no oscillations in NP1227 class1, which could explain why inactivation of NP1227 class1 does not attenuate the oscillations in PNs.

### 2.4 Temporal dynamics of firing in MBON-*α*1

Figure 5a shows the responses of the somatic membrane potential of MBON-*α*1 *in vivo* (red) and *in silico* (blue) before and after olfactory associative learning. The solid plots represent the values of the somatic membrane potential, and the black dots above them represent the detected spike timing. The gray arrows indicate the onset of 3-octanol input. 3-octanol was given for one second. We calculated the firing frequency for each 50 ms time window from the spikes and plotted the average firing frequency transition over five trials as dotted curves. The procedures for detecting spikes and calculating their frequency are described in Supplementary Note 2.

**Figure 5:**
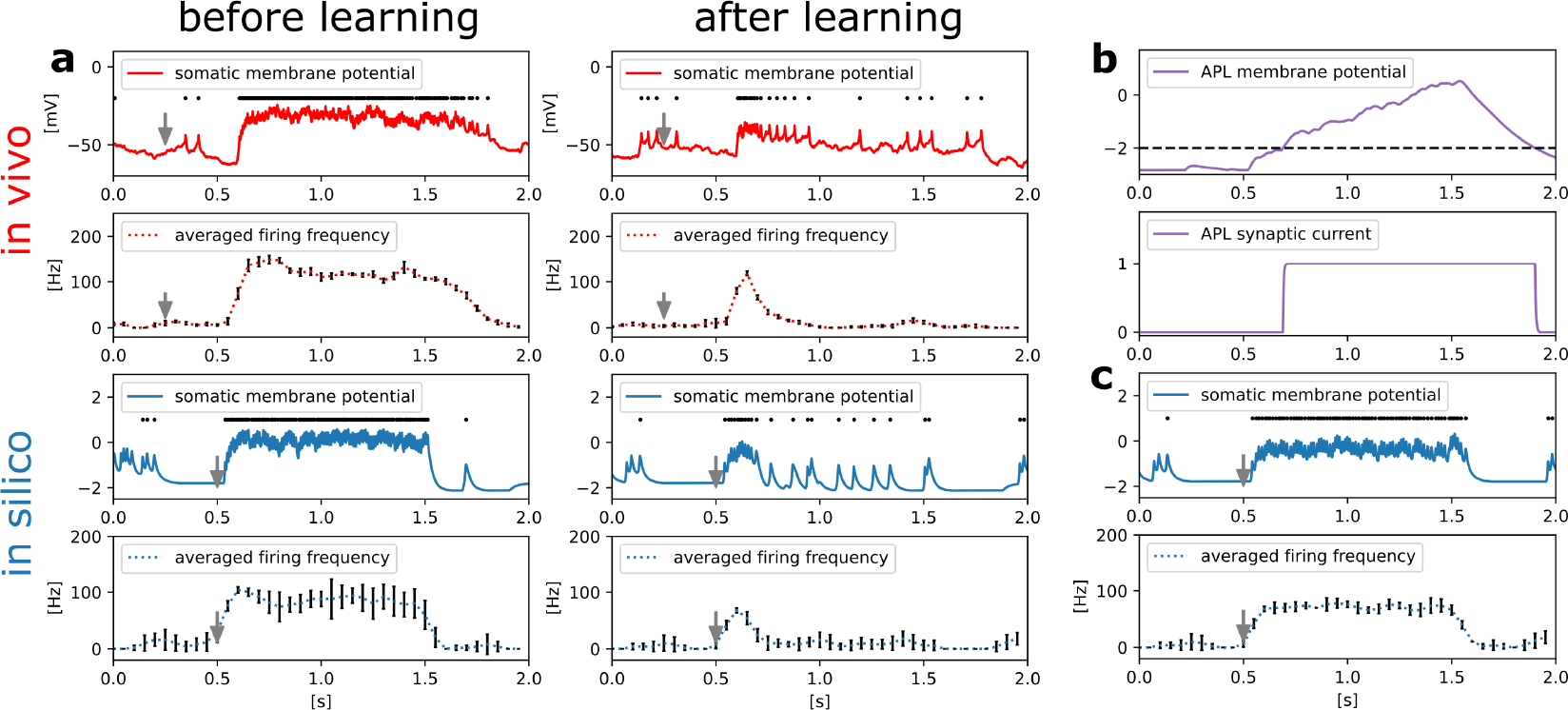
Responses of MBON-*α*1. **a** Comparison of somatic membrane potential of MBON-*α*1 *in vivo* (red) and *in silico* (blue) before and after olfactory associative learning. The gray arrows indicate the onset of a one-second-long application of 3-octanol. *In silico*, the odor input caused ORNs to fire instantaneously, whereas *in vivo*, there was a delay for the odor to travel through the tubing and reach ORNs. Example responses of the somatic membrane potential for a single trial (solid line) and the temporal dynamics of firing frequency averaged over five trials dotted line). Error bars represent standard deviation over five trials. The black dots above the solid lines represent spikes. **b** The membrane potential and synaptic current of APL are shown. The black dotted line represents the threshold for the release of the synaptic current of APL. **c** Example responses of the somatic membrane potential of MBON-*α*1 in the absence of APL for a single trial (solid line) and its averaged firing frequency over five trials (dotted line). Error bars represent standard deviation over five trials. The black dots above the solid line represent spikes.

*In vivo*, whereas odor-evoked firing frequency of MBON-*α*1 is constantly high before learning, it decreases rapidly after learning. We reproduced this characteristic temporal dynamics of firing *in silico*. The somatic membrane potential and synaptic current of APL are shown in Figure 5b to illustrate how the temporal dynamics occur *in silico*. While MBON-*α*1 fires immediately after odor onset, the membrane potential of APL reaches the threshold with a delay due to its slow neuronal dynamics. This delayed inhibition from APL may contribute to suppressing the firing of MBON-*α*1 from approximately *t* = 0.7, together with LTD at KC*>*MBON-*α*1 synapses. We tested this possibility in silico by examining the activity of MBON-*α*1 while inactivating APL (Fig. 5c). Without APL, the odor-evoked firing frequency of MBON-*α*1 is constantly high even after learning, and this result indicates that APL is essential for the temporal activity of MBON-*α*1.

### 2.5 FPGA Implementation

We implemented the network model on a Xilinx Artix-7 XC7A35T FPGA on a Digilent cmod-a7 board using Xilinx Vivado 2016.4. Figure 6 presents an overview of the implementation. As the network has nine types of neurons, namely four LN subclasses, PN, KC, APL, MBON, and SMP354 neuron, we constructed nine PQN engines corresponding to each of them. The weights of synaptic connections, input current, synaptic current, and neuronal internal state variables are stored in block RAMs. Spike signals of ORNs were generated by the PC and sent to the FPGA through a serial communication bus. The spike signals were composed of 11 bits representing the indices of ORNs, which were initially stored in the FIFO buffer; the synaptic currents of ORNs were calculated using the SC engine. The accumulators calculated the input currents for neurons from the synaptic currents in parallel. The antennal lobe and the mushroom body are distant, and only PNs provide a one-way connection from the antennal lobe to the mushroom body. Therefore, we built three accumulator blocks, a, b, and c, which consisted of seven, three, and one accumulator(s), that were responsible for the processing inside the antennal lobe, between the antennal lobe and the mushroom body, and inside the mushroom body, respectively. The weights *w* of the KC*>*MBON-*α*1 synapses are represented by two bits to realize the LTD, whereas all other synaptic weights are represented by one bit. The PQN controller activates each PQN engine in turn. Each PQN engine receives the current values of the internal variables, input currents, and synaptic currents of the corresponding type of neurons. It then returns the next step values of the internal variables and synaptic currents. The details of the PQN engines are provided in the Methods section. A reward signal was also transmitted using serial communication to the LTD unit, which triggered the LTD of KC*>*MBON-*α*1 synapses. The LTD unit holds the indices of KCs that have fired in the previous five seconds, and when the reward signal arrives, it rewrites *w* of synapses made by those KCs onto MBON-*α*1.

**Figure 6:**
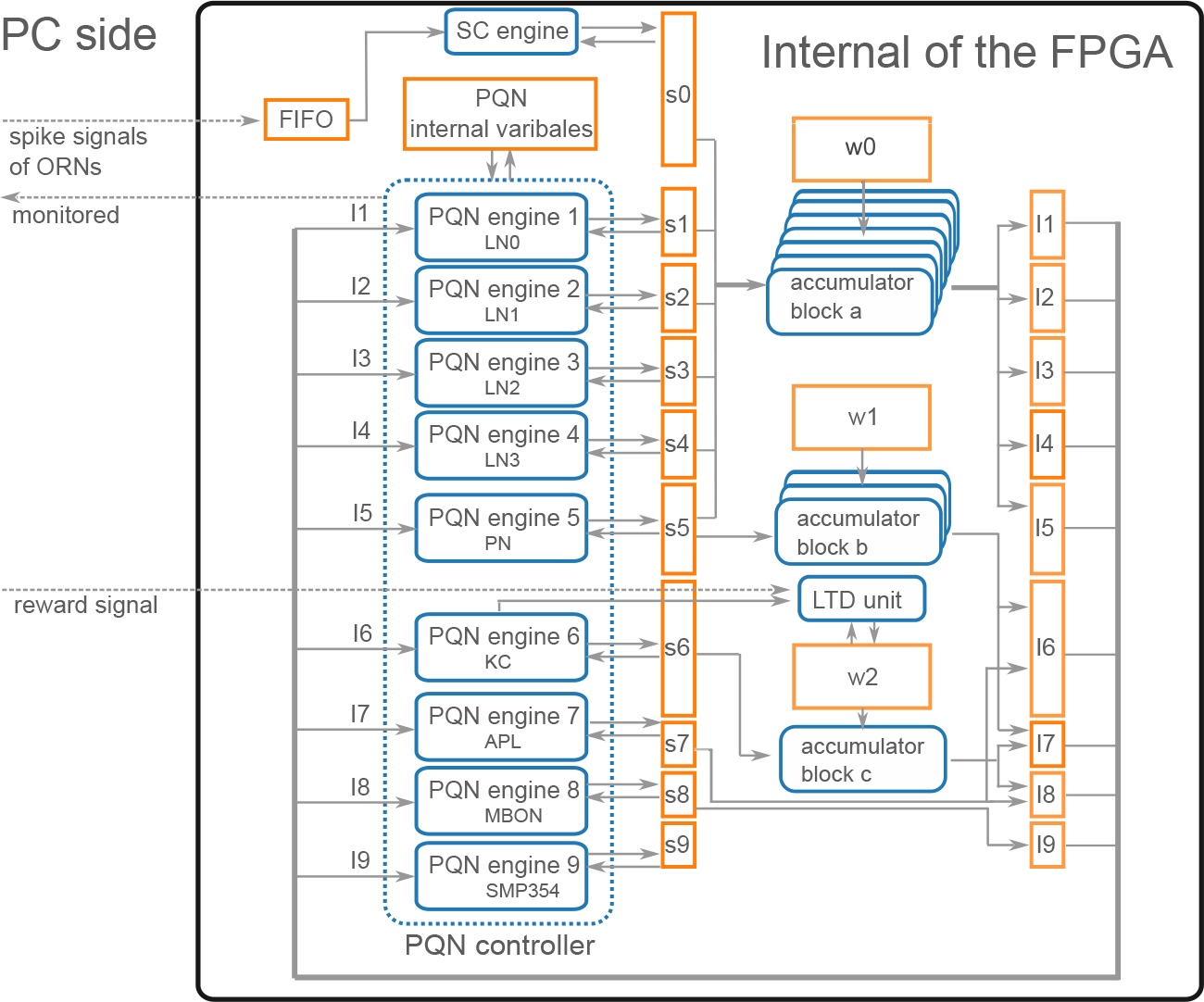
Architecture overview. The rectangle, rounded rectangle, and arrows represent the block RAM, computation unit, and data flow, respectively. PQN engines, where *x* ranges from 1 to 9, simulate the activities of each type of neuron. LN0, LN1, LN2, and LN3 correspond to Krasavietz class1, Krasavietz class2, NP1227 class1, and NP2426 class1, respectively. s*x*, where *x* ranges from 0 to 9, and I*y*, where *y* ranges from 1 to 9, are the synaptic and input currents, respectively. w0, w1, and w2 store the weight of synaptic connections. The PQN internal variables store neuronal internal state variables. Spike signals of ORNs are sent from the PC using serial communication. They are temporarily stored in the FIFO and subsequently converted to synaptic currents using the SC engine. The accumulator blocks a, b, and c comprise seven, three, and one accumulator(s), respectively, and each accumulator calculates the input currents from the synaptic currents in parallel. The PQN controller activates each PQN engine in turn to simulate neurons. Each PQN engine receives the internal variables, input currents, and synaptic currents of the corresponding type of neurons and returns the next step values of the internal variables and synaptic currents. The reward signal from the PC activates the LTD unit and reduces the weights of KC*>*MBON-*α*1 synapses that are stored in part of w2. The information required for each result section, such as the spike information of the PQN neurons, values of membrane potential, and synaptic currents, is selected and sent to the PC and stored.

Figure 7a shows the resource consumption of this implementation. The look-up tables (LUTs) are truth tables that were used primarily for addition calculations in this implementation. Digital signal processors (DSPs) are blocks for complex calculations that were used to multiply the state variables. Flip-flops (FFs) and block random-access memories (BRAMs) are memory elements. Most BRAMs store synaptic weights, whereas the rest store state variables. A mixed-mode clock manager (MMCM) was used to generate a 100 MHz clock. Figure 7b lists the on-chip power consumption of each resource estimated by Vivado. The static represents the steady-state leakage power of the device and is independent of the circuit design. The total power consumption is approximately 0.36 watts.

**Figure 7:**
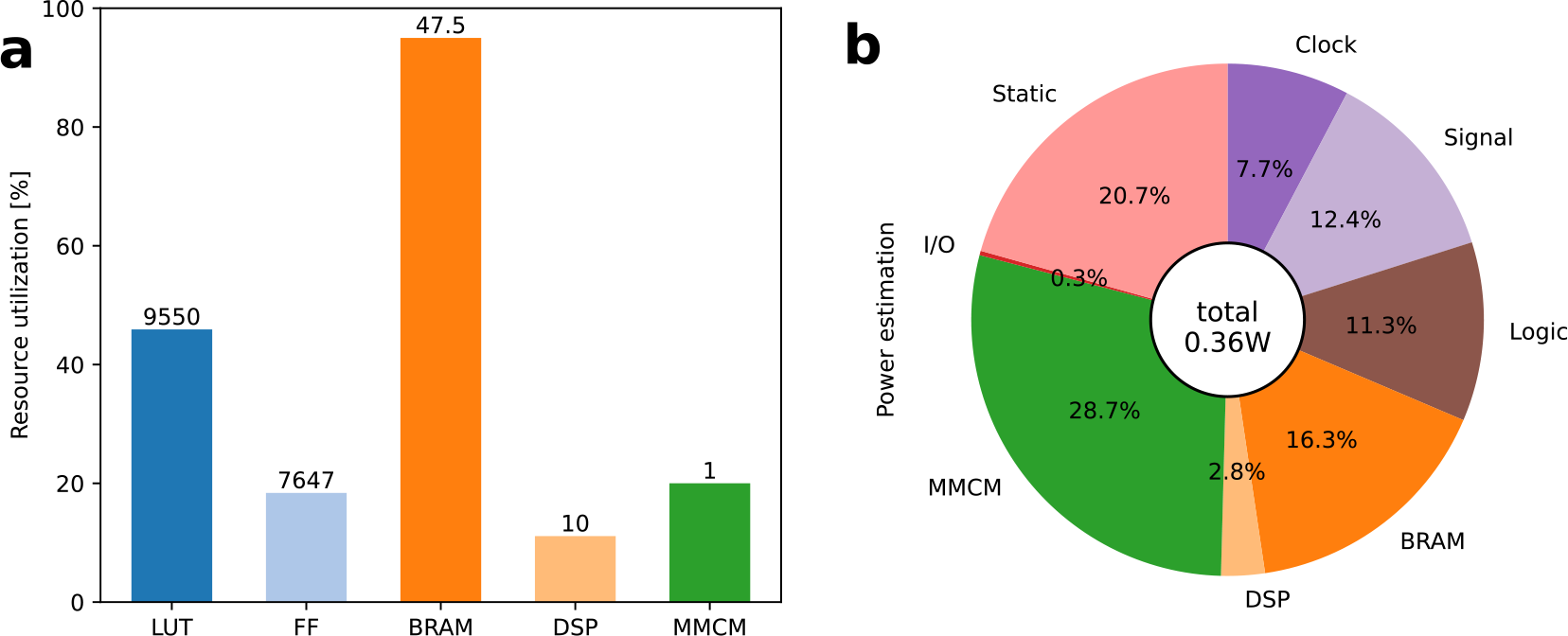
Results of FPGA implementation. **a** Utilization ratio for each type of resource. The numbers above the bar indicate the number of units used. **b** Power estimation for each type of resource.

## 3 Discussion

In this study, we built the first data-driven SNN model of the olfactory nervous system of *Drosophila melanogaster*. Our modeling approach proposed a way to overcome the trade-off between replicating the detailed biological data (the connectome and electrophysiological activities of neurons) and the computational cost, such that the model can run in real-time on a low-power SiNN chip while reproducing the characteristic neuronal activities in the brain. Features of previous data-driven models [8][9][10] that reproduced parts of the mammalian cortex and hippocampus as well as this work are compared in Table 1. Specifically, our model went beyond the preceding models in the following four aspects: the higher reproducibility of (1) synaptic connectivity, (2) characteristic spiking activities, (3) neuronal functions, and (4) the lower computational cost. Whereas the preceding models reproduced the electrophysiological and morphological properties of each type of neuron using multicompartmental ionic-conductance-based models, our model reproduced electrophysiological properties using the PQN model, which requires a lower computational cost. In [8][10], the Tsodyks–Markram (TM) synapse model [43] with a stochastic mechanism was used to accurately reproduce synaptic physiology, whereas in [9], the double exponential synapse model reproduced the rising and decaying time constants of the synaptic current for each type of synaptic connection. In this study, the decay time constant of the double exponential synapse model was fitted to electrophysiological data for the corresponding type of neurotransmitter. In the preceding models, synaptic connections were randomly determined based on the position and morphology of individual neurons and statistical information for each neuron type. However, in this model, they were based on the connectome [21][37] identified from comprehensive electron microscopy images. In the preceding models, the vast number of neurons and complex structures of the mammalian brain limited the validation of the models. In [8] and [9], synchronous oscillations at the network level were validated, but not for each type of neuron. Spiking activities were not examined in [10]. Additionally, the preceding models did not reproduce the function of the network, as mammalian cortical and hippocampal functions at the circuit level have not yet been elucidated. In contrast, because the olfactory nervous system has a smaller network size and its function is clearer, we were able to demonstrate that our model successfully reproduces olfactory associative learning, characteristic spiking activities of each type of neuron, such as odor-evoked oscillatory firing in PNs and LNs, absence of oscillations in KCs, different contributions of LN subclasses to the formation of oscillations, and temporal dynamics of firing in MBON-*α*1. Whereas the preceding models required supercomputers owing to their enormous computational cost, our model was light enough to be simulated on an entry-level low-cost FPGA chip at 0.36 watts, which may be acceptable for small robots and portable AI devices. In addition, whereas the simulation speed in [9] was approximately 1600 times slower than real time, our model performs real-time simulations.

**Table 1:** Comparison of the data-driven SNN models.

|  | This paper | [8] | [9] | [10] |
| --- | --- | --- | --- | --- |
| Target | *Drosophila* olfactory nervous system | microcircuit of rat neocortex | CA1 of rat hippocampus | CA1 of rat hippocampus |
| Model | PQN model | ionic-conductance -based model | ionic-conductance -based model | ionic-conductance -based model |
| Scale | 2200 neurons | 31000 neurons | 340000 neurons | 400000 neurons |
| Reproducibility of neuronal electrophysiology | high | high | high | high |
| Reproducibility of neuronal morphology | no | high | high | high |
| Reproducibility of synaptic properties | low (PQN synapse) | high (TM with stochastic) | medium (double exponential) | high (TM with stochastic) |
| Reproducibility of synaptic connectivities | high (connectome-based) | medium (morphology-based statistical method) | medium (neural distance -based statistical method) | medium (morphology-based statistical method) |
| Reproducibility of characteristic spiking activities | high | medium | medium | low |
| Reproducibility of functions | high | no | no | no |
| Computing environment | FPGA (0.3W) | Supercomputer (4-rack IBM Blue Gene/Q) | Supercomputer (3488 processors) | supercomputer |
| Simulation speed | real time | not available | 1642 times slower | not available |

There also are differences between our model and the latest preceding model [26] of the *Drosophila* olfactory system. Unlike our model, the preceding model is not data-driven. Whereas the preceding model stochastically determines the connections between layers such as ORN*>*PN and PN*>*KC, our model precisely reproduces the connections based on the connectome database. As for the target network components, whereas the preceding model consists of PNs, LNs, KCs, APL, and MBON, our model has another MBON and SMP354 neuron in addition, reproducing the valence-balance model [44][36], where learning-induced plasticity in the KC*>*MBON synapses tips the balance of valence signals of MBONs. This competitive memory circuitry is important because it is the basis for the interactions among MBONs that are responsible for flexible and complex behavioral decisions associated with memory. As for the spiking dynamics, the characteristic spiking activities of each neuron are not considered in the preceding model.

For example, the spiking activities of PNs and LNs are not calculated by spiking neuron models but are generated by the Poisson process. The activities of MBONs are represented using nonlinear activation functions. KCs are described by the LIF model, and their firing properties are not fitted to the *in vivo* data.

The peak frequency of PN oscillation in this model was approximately 24 Hz, whereas experimentally observed peak frequency in the antennal lobe was 10–15 Hz [27]. In the antennal lobe, PNs and LNs are connected via glomeruli, which are neuropils comprising the dendrites and axons of PNs, LNs, and ORNs. However, the model does not consider the dynamics of the glomeruli, which may cause a gap in peak frequencies. In addition, the proportion and detailed connections of the four subclasses of LNs are not known; therefore, they were not incorporated into the model and may have affected the peak frequency. A more detailed model awaits to be built to clarify the mechanism and function of oscillations in the antennal lobe.

To examine the oscillations (Fig. 4), we only applied 3-octanol to the network. This is because the magnitude of PN’s oscillations greatly depends on the identity of odors both *in vivo* [27] and *in silico* (Supplementary Figure 9). Since our intention was to measure the effect of inactivation of the LN subclass on PN’s oscillations, we used only one type of odor. In the future, we will comprehensively examine the relationship between oscillations and odors and clarify why the magnitude of the oscillations differs between odors.

In honeybees, oscillations in the antennal lobe are necessary for distinguishing between similar odors [41]. In locusts, oscillations appear not only in the antennal lobe but also in KCs, and they are believed to contribute to sparse representation of odors in the KC population [42]. Although the role of oscillations in *Drosophila* remains unclear, oscillations likely contribute to the processing of odor information given the similarity of olfactory network structure between different insects. One possible candidate is the generation of sparse representation of odors in the antennal lobe.

In this study, the PQN model employs function *m*(*I*), which was not incorporated into the original PQN model [16]. This function performs a nonlinear transformation of the stimulus input so that the membrane potential behaves as expected in response to a wide range of stimulus inputs. However, this function does slightly complicate the model and has no biological counterpart. By changing the parameters and adjusting the dynamics, we expect to be able to remove this function in future works.

As shown in Figure 5, after olfactory associative learning, MBON-*α*1 fires for approximately 250 ms, immediately after the arrival of the odor signal, and subsequently enters a resting period, successfully reproducing the temporal firing observed in [29] in MBON-*γ*1pedc. To our knowledge, there has been no report on the mechanism underlying this firing dynamics characteristic for the post-learning response. The result of our simulation suggests that the delayed activation of APL contributes to shaping this activity pattern. Thus, our modeling not only reproduces observed physiological data but also provides mechanistic insight by proposing an experimentally testable hypothesis.

The SiNN implemented in this study operates at the same speed as the olfactory nervous system with a 100 MHz clock signal. However, if we use a higher clock, the model can provide accelerated simulations, albeit with increased power consumption. For example, we confirmed that the model can simulate four times faster than real-time using a 400 MHz clock with a Xilinx Virtex UltraScale+ xcvu37p-fsvh2892-3-e FPGA. In this implementation, the estimated power consumption was about 4W. The power efficiency and simulation speed can be further improved by using Application Specific Integrated Circuits (ASICs).

As shown in Figure 7b, most of the power is consumed by the MMCM, BRAMs, and the steady-state leakage (Static). Except for a few BRAMs that are used to store the neuronal state variables, these resources are not directly used to compute the neuronal dynamics. Ignoring the reproducibility of the spiking properties and using I&F-based models instead of PQN might reduce the power of clocks, signals, logic, and DSPs. However, these resources consume only 18.5% of the total power and their impact on the overall system is expected to be small. Ionic-conductance-based models can reproduce the dynamics of the spiking process as accurately as or better than the PQN model. However, they have many exponential terms that consume a large number of DSPs in FPGA implementations [45][46]. Even in the most well-optimized implementation [46], it requires more than 20,000 LUT units and more than 100 DSPs to build a network of 2,000 neurons, which would lead to significantly higher power consumption.

Our modeling approach is applicable to not only FPGAs but also ASICs. Conversion from FPGA to ASIC improves power efficiency by a factor of 14 to 20 [47][48]. The network reproduced in this study accounts for approximately 2% of the entire brain. Thus, our approach enables the construction of an ASIC chip that simulates the entire *Drosophila* brain while consuming approximately 1 watt. Such chips have considerable potential in the engineering and scientific fields. Because of its low power consumption, the chip can be mounted on small insect-like robots. The resulting system is expected to move around autonomously, solve unknown tasks, and adapt to changes in the environment, similar to insects. In addition, owing to its intrinsic power efficiency, the chip can serve as a sufficiently fast simulator of the whole brain within the constraints of the power supply typically available in laboratories. It can facilitate long-term measurement of neuronal activities and is expected to contribute to the analysis of phenomena with long timescales, such as continuous learning and forgetting.

## 4 Methods

### 4.1 Details of the network

Synaptic connections were determined using the connectome database hemibrain v1.2.1 [21][37]. Since this model targets the olfactory nervous system of the right hemisphere, we first obtained the number of neurons of each type in the right hemisphere from the hemibrain server using the NC function of the neuprint-python library. We then determined the connections between neurons using the fetch neurons function of the neuprint-python that returns the number of synaptic connections between neurons. Connections with more than ten synapses were assumed to have sufficient strength and their weight *w* was set to 1. Otherwise, *w* was set to 0. Note that the connections of LNs were determined based on a previous study [32].

A variety of LN subclasses were reported [38][32], and [32] identified four subclasses, each with different spiking properties. However, the connectome database [21][37] does not describe which subclass each LN belongs to. Therefore, the connection of LNs was determined based on [32] where the probabilities that each LN subclass has a connection to a certain glomerulus were shown. In the antennal lobe, glomeruli are neuropils comprising axons and dendrites of PNs, LNs, and ORNs. ORNs and PNs are generally connected to only one glomerulus. We first determined the subclasses to which 191 LNs belong. As the proportion of each subclass is unknown, we set the number of NP2426 class1 to 47 and the remaining to 48 to ensure that the distribution of subclasses was as even as possible. Next, for each LN, we randomly determined whether each LN innervates each glomerulus according to the innervation probabilities shown in [32]. If an ORN/PN and LN innervate the same glomerulus, the ORN/PN was assumed to make a synaptic connection onto the LN, and the synaptic weight *w* was set to 1. For example, LNs NP1227 class1 connect to glomerulus DA1 with a probability of 75% [32]. Based on this probability, we determined whether each LN NP1227 class1 connects to glomerulus DA1.

Each ORN expresses one of the olfactory receptors, each of which has different odor selectivity. A previous study [49] described a correspondence table between glomerulus and OR types. We used this table and the glomerulus type for each ORN listed in the connectome database to determine the OR type for each ORN. When multiple OR types were assigned to a single glomerulus type, one OR was randomly selected.

Figure 8(a) shows part of the connection structure from ORNs to PNs in the model. The black squares represent the presence of connections. On average, each ORN projects to 1.6 PNs, and each PN receives input from 24.0 ORNs. ORNs and PNs were sorted based on their glomerulus type, the borders of which are represented by blue lines. ORNs generally project to all the PNs in the same glomerulus type [50]. This convergent projection is considered [51] to enable PNs to produce reliable output by averaging the input from a large number of ORNs, whereas the responses of ORNs to odors are noisy and unreliable [34]. LNs receive inputs from a wide range of ORNs and extensively inhibit PNs and LNs. On average, each LN receives input from 1337.4 ORNs and inhibits 90.9 PNs. LNs are considered to contribute to the gain control of the input from ORNs [52] and to the generation of oscillations in the antennal lobe [27].

**Figure 8:**
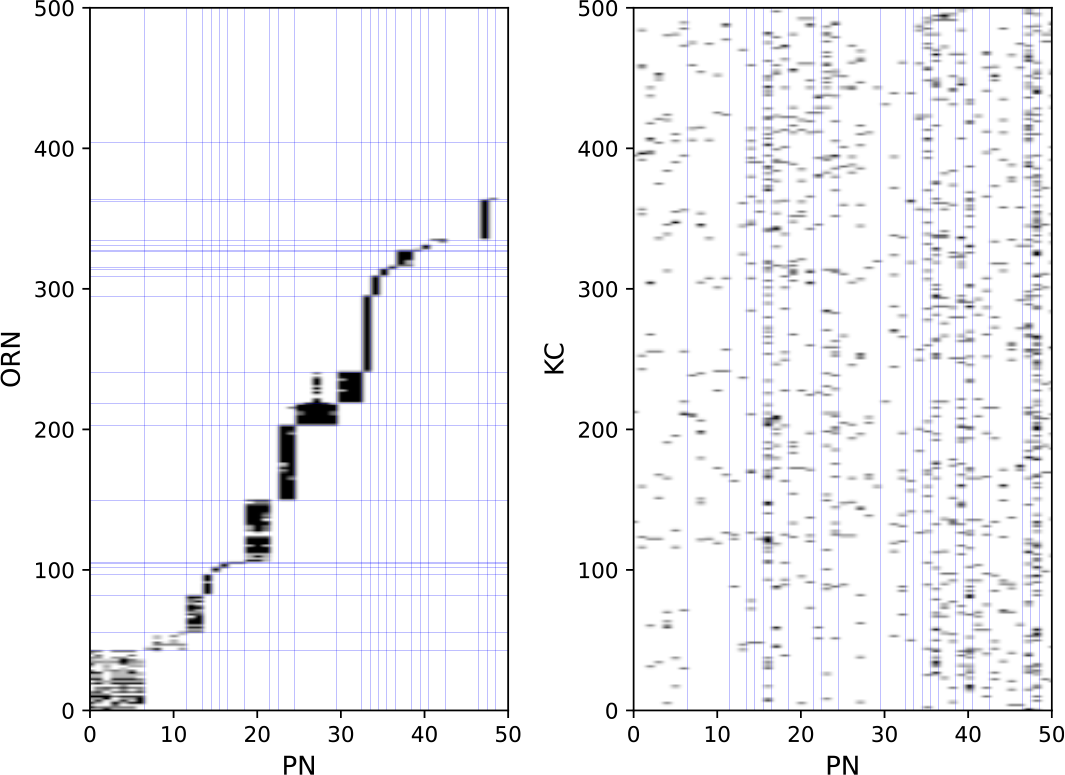
Synaptic connections from ORNs to PNs (a) and PNs to KCs (b).

PNs extend their axons to the entrance of the mushroom body, where they provide excitatory input to KCs. On average, each PN projects to 67.6 KCs, and each KC receives input from 4.2 PNs. Figure 8(b) shows part of the connection structure from PNs to KCs in the model. In contrast to the connections between ORNs and PNs, there is no regularity in the connections between PNs and KCs, which was confirmed in a previous study [53]. APL receives input from almost all KCs and PNs and returns inhibitory feedback to KCs and MBONs.

There are 28 MBONs, or 44 including atypical MBONs [54], at least some of which signal either positive or negative valence [55]. While how MBON signals are further processed by the downstream circuits to determine the behavioral output is still largely elusive, the connectome study discovered that postsynaptic neurons of the MBONs typically receive synaptic input from more than one type of MBONs [54], suggesting that valence signals could be integrated by those neurons. A recent study characterized such a circuit motif experimentally [36]. A cluster of 8-10 neurons named UpWind Neurons (UpWiNs) directly and indirectly integrates excitatory and inhibitory input from MBON-*α*3 and MBON-*α*1, respectively. Direct optogenetic activation of MBON-*α*3 induces upwind locomotion, which can be interpreted as an olfactory approach behavior [56], while activation of MBON-*α*1 does not induce such behavior [36]. Experiments using compartment-specific optogenetic activation of dopaminergic neurons demonstrated that *α*3 and *α*1 are an aversive- and appetitive-memory compartment, respectively [57]. Moreover, optogenetic activation of UpWiNs triggers robust upwind locomotion [36]. Thus, the UpWiN cluster is one of the sites where signals of opposite memory valence are integrated and translated into olfactory navigation behavior. Since neurons in the UpWiN cluster are heterogeneous in their anatomy and connectivity, in our model, we focused on one of the neurons, SMP354, (bodyId in hemibrain is 390003153), which receives direct synaptic input from both MBON-*α*3 and MBON-*α*1. Because there is no specific genetic driver to label this particular neuron, we were unable to use experimentally determined electrophysiological parameters for this neuron. Although our model is simplified in terms of the readout mechanism of the mushroom body signals, we believe that the SMP354 circuit represents one of the common motifs that interpret the population signals of MBONs.

If a reward or punishment is given to a fly with an odor, the fly will learn to approach or avoid that odor thereafter [58]. Multiple studies [29][59][60] indicate that this olfactory associative learning is caused by long-term depression of KC*>*MBON synapses. In the case of the circuit associated with UpWiNs, it has been experimentally demonstrated that induction of plasticity in *α*1, which mimics appetitive conditioning, depresses olfactory responses of MBON-*α*1. This in turn potentiates responses of UpWiNs, whose naïve odor responses are typically weak [36]. In this model, when a reward stimulus was given, the weights of KC*>*MBON-*α*1 synapses were weakened if the presynaptic KCs had been firing within five seconds. Although the magnitude of the decrease in the synaptic weight after a single learning is not clear, we set the initial value of *w* to 1 and the weakened value to 0.25. Reward stimuli are transmitted to KC*>*MBON synapses through dopaminergic neurons innervating the mushroom body [55]; however, this pathway was not modeled in this study.

### 4.2 Electrophysiological measurements

#### 4.2.1 Recording from PNs

Whole-cell patch-clamp recordings from PN somata were performed as previously described [33]. Briefly, the brain of *w;UAS-ReaChR::Citrine(attP40)/+;VT033006-Gal4(attP2)/+* female flies [61][62], 3 days post eclosion, was removed from the head capsule and fixed on a glass slide with surgical glue (GLUture, Abbott). Part of the perineural sheath covering the antennal lobe was removed to obtain an access to cell bodies. The external saline added on top of the plate was circulated throughout the experiment. A patch pipette was pulled from a thin-wall glass capillary (1.5 mm o.d./ 1.12 mm i.d., TW150F-3, World Precision Instruments). Resistance of the pipette was typically 8-10 MΩ. The internal solution contained (in mM) 140 KOH, 140 aspartic acid, 10 HEPES, 1 EGTA, 4 MgATP, 0.5 Na3GTP, 1 KCl, and 13 biocytin hydrazide (pH ∼7.2, osmolarity adjusted to ∼265 mOsm). Electrophysiological recordings were made with a Multiclamp 700B amplifier (Molecular Devices) equipped with a CV-7B headstage. Signals were low-pass filtered at 2 kHz and digitized at 10 kHz. Multiple levels of depolarizing currents were injected into the soma of individual PNs to examine the relationship between the input current and the membrane potential or spike output. PNs were identified based on the signals from Citrine as well as biocytin included in the internal solution.

#### 4.2.2 Recording from MBONs

*In vivo* whole-cell current-clamp recordings from MBON-*α*1 and optogenetic trainings were performed as previously described [29][36]. Female flies with the genotype of *10xUAS-ChrimsonR-mVenus (attP18)/w; R71C03-LexA (attP40)/LexAop-GFP (attp5); MB043C/+* reared on conventional cornmeal-based food were collected on the day of eclosion, transferred to all-transretinal food (0.5 mM) and kept in the dark for 48-72 hr until experiments. The patch pipettes were pulled for a resistance of 4-6MΩ and filled with pipette solution containing (in mM): L-potassium aspartate, 140; HEPES, 10; EGTA, 1.1; CaCl2, 0.1; Mg-ATP, 4; Na-GTP, 0.5 with pH adjusted to 7.3 with KOH (265 mOsm). The preparation was continuously perfused with saline containing (in mM): NaCl, 103; KCl, 3; CaCl2, 1.5; MgCl2, 4; NaHCO3, 26; N-tris(hydroxymethyl) methyl-2-aminoethane-sulfonic acid, 5; NaH2PO4, 1; trehalose, 10; glucose, 10 (pH 7.3 when bubbled with 95% O2 and 5% CO2, 275 mOsm). Whole-cell recordings were made using the Axon MultiClamp 700B amplifier (Molecular Devices). MBON-*α*1 was visually targeted by the GFP signal with a −60X water-immersion objective attached to an upright microscope. Cells were held at around 60 mV by injecting hyperpolarizing current. Signals were low-pass filtered at 5 kHz and digitized at 10 kHz. Data acquisition and analyses were done by custom scripts in MATLAB (MathWorks). 3-octanol (OCT) and 4-methylcyclohexanol (MCH) were presented to flies with custom odor delivery system after diluting to 1% of the saturated vapors. After recording baseline responses by alternately presenting OCT and MCH five times (duration, 1 s; interval, 30 s), OCT was paired with 625 nm LED photostimulation (pulse duration, 1 s; frequency, 0.5 Hz; power, 17 mW/mm2) for 1 min. MCH was presented without photostimulation for 1 min. 1.5 min later, post-pairing responses to both odors were recorded five times. The pairing resulted in selective depression of OCT responses, which is consistent with a previous study [36]. The I-V relationship was measured before pairing by injecting 1-s square pulses with incrementing amplitudes (0–10 pA, 2 pA steps).

### 4.3 ORN input data

The input data were generated using the DoOR dataset [49], which comprehensively reports the response properties of ORs of *Drosophila*. The dataset shows the response intensities of each OR for a wide variety of odorants. Given a certain odor, the firing frequency *r* of an ORN that expresses a certain OR is given by

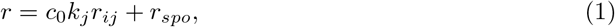

where *r*_*ij*_ is the response intensity of the *i*th OR to the *j*th odorant, and its value ranges from 0 to 1. *r*_*spo*_ represents the spontaneous firing frequency, which was set to 8 from the average value examined in [63]. *k*_*j*_ is a constant that abstractly refers to the concentration of the *j*th odorant; its values range from 0 to 1 and are listed in Supplementary Note 4. As ORNs fire at approximately 200 Hz in response to the most favorable odorants [64], parameter *c* was set to 192, such that the maximum firing frequency *r* would be 200 when *r*_*ij*_ and *k*_*j*_ were 1. Based on the Poisson process, each ORN generates a spike with probability *rdt* at every time step, where time step *dt* is 1 ms. In the input dataset, six odorants were applied sequentially for one second every five seconds. The synaptic currents from ORNs were calculated using the following equation:

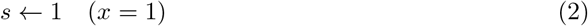

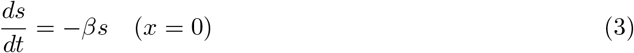

where *x* represents the spiking information of an ORN. *x* is 1 when a spike is emitted in the current time step by an ORN and 0 otherwise. ORNs are cholinergic [65], and their *β* was set to 203.125 as well as the other synapses.

We prepared three types of input data for the *in silico* experiments. In the first type of the input data, one of the six odorants, 3-octanol, cis-3-hexenol, cyclohexanone, 2,3-butanedione, 2-hexanol, and ethyl butyrate, was applied in turn for one second every five seconds. In the second type of data, 3-octanol, was applied for ten seconds every twenty seconds. In the third type of data, the same six odorants as the first type were applied in turn for ten seconds every twenty seconds. In all *in silico* experiments, the first type of data was initially given for 300 seconds, during which time the PN’s homeostasis was adjusted (details are described in Supplementary Note 3). Subsequently, the first type of data was continuously provided, and experiments on associative learning and the activity of MBON-*α*1 were conducted. In contrast, in the experiments on the oscillations in the antennal lobe, the second type of data was applied following the 300-second homeostatic period. The third type of data was only used in the experiment to show the variations in oscillations for each odor (Supplementary Figure 9).

### 4.4 Calculation of the power spectra

The fft package of the NumPy module in Python was used to calculate the power spectra of the virtual LFP. Here, only 8 seconds of the 10-second responses of the virtual LFPs to the odor, excluding the first and last seconds, were used to compute the power spectra.

### 4.5 PQN model

The piecewise quadratic neuron (PQN) model [11][12][13][14][15][16] is a qualitative neuron model designed to replicate a wide variety of neurons in the nervous system and to be efficiently implemented on digital arithmetic circuits. Compared with other qualitative models [66][67][68], the PQN model possesses additional parameters, enabling it to represent more functional forms and reproduce a variety of neurons, each with its unique dynamical structure. In addition, although other qualitative models have cubed variable terms, which consume a vast amount of circuit resources in digital arithmetic circuits, the PQN model uses piecewise functions composed of a squared term to represent comparable dynamics and consumes few circuit resources.

The nervous system of *Drosophila* primarily comprises unipolar neurons, the soma of which is separated from the rest of the cell by a long and thin membrane. In the patch-clamp recording from the soma, only action potentials with extremely small amplitudes were observed. This is attributed to the fact that the action potentials are generated in the axon and propagated with decay to the cell body [69]. Therefore, we modeled PNs, KCs, and MBONs using two-compartment models; one compartment corresponded to the soma, and the other contained axons and dendrites. In contrast, in the soma of LNs, sufficiently large action potentials were observed [32]; therefore, they were modeled using single-compartment models. APL is an inhibitory, non-spiking neuron whose axons extend to the whole mushroom body. It was reported [33][70] that APL performs local inhibition; however, the details are not clear. Therefore, in this study, we modeled APL as a simple non-spiking neuron with a single-compartment.

The equations of the PQN model in the single-compartment version for LNs, APL, and SMP354 are given by

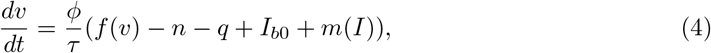

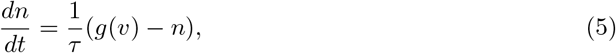

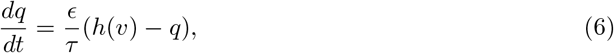

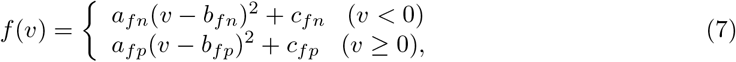

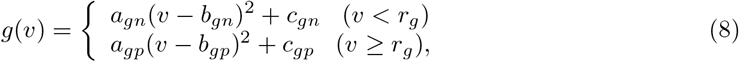

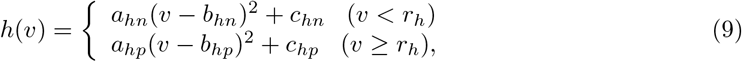

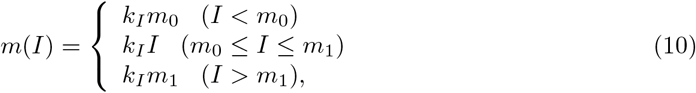

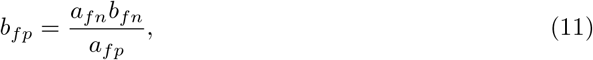

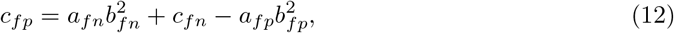

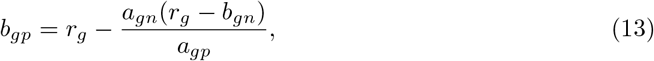

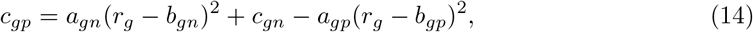

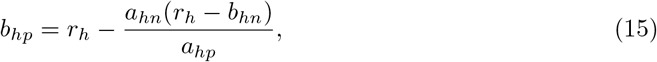

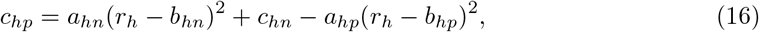

where *v, n*, and *q* correspond to the membrane potential, recovery variable, and slow variable, respectively. Parameter *I*_*b*0_ is a bias constant. Parameter *I* represents the stimulus current. The function *m* performs a nonlinear transformation of *I*, adjusting the scale of *I* with parameter *k*_*I*_ and extending the dynamic range with parameters *m*_0_ and *m*_1_. Synaptic currents from other neurons and current injections shown in Figure 2 were given to *I*. The parameters *τ*, *ϕ*, and *ϵ* determine the time constants of the variables. The parameters *r*_*g*_, *r*_*h*_, *a*_*x*_, *b*_*x*_, and *c*_*x*_, where *x* is *fn, fp, gn, gp, hn*, or *hp*, are constants that determine the nullclines of the variables. The parameters *b*_*fp*_, *c*_*fp*_, *b*_*gp*_, *c*_*gp*_, *b*_*hp*_, and *c*_*hp*_ are determined by other parameters such that the nullclines are continuous and smooth. All variables and parameters are purely abstract with no physical units. The initial values of all state variables were set to zero.

The equations of the two-compartment version for KCs and MBONs are given by

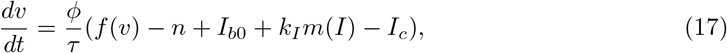

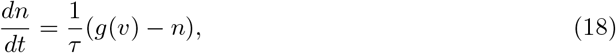

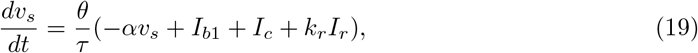

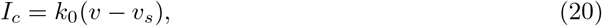

where *v* and *n* are the membrane potential and recovery variables in the axonal compartment, respectively, and *v*_*s*_ is the membrane potential of the somatic compartment. Parameters *θ, α*, and *I*_*b*1_ are the time constant, bias constant, and leakage constant, respectively. *I*_*c*_ represents the internal current that flows from the axonal compartment to the somatic compartment, and *k*_0_ is its kinetic parameter. When synaptic currents are given to *I*, the current injected into the soma (Fig. 2) is given to *I*_*r*_, and *k*_*r*_ is its scaling parameter.

In PNs, homeostatic control of synaptic efficacy has been indicated [65]. Although various types of homeostatic mechanisms are found in neurons, we modified the equation for PNs based on the mechanism of synaptic scaling proposed in a previous study [71], where the weights of synaptic connections were gradually scaled according to the activity level of the postsynaptic neuron. The equations used are as follows:

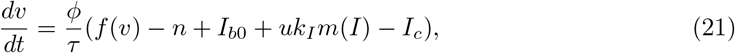

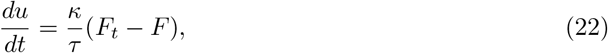

where *F* and *F*_*t*_ represent the neuronal current firing frequency and target firing frequency, respectively. The equation of *I*_*c*_ and the differential equations of *n* and *v*_*s*_ are the same as those in KCs and MBONs (Eqs. (18) and (20)). The parameter *κ* determines the time constant. Note that the value of *u* is fixed between 0 and 1.

The synaptic current is calculated as follows:

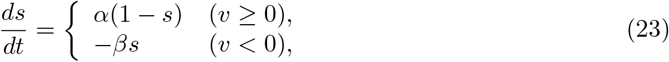

where *s* denotes the synaptic current and the parameters *α* and *β* determine the time constants. This synaptic model is a qualitative version of the simplified kinetic model of chemical synapses [72]. Although the dynamics of each synapse are unclear, the decay constants of cholinergic synapses from PN to KC and GABAergic synapses in cultured embryonic neurons have been investigated [73][74] and are both approximately 5 ms. Therefore, the values of *β* were set to 203.125 so that their decay time constants were close to 5 ms. The value of *α* was chosen to be 250 so that a single spike results in a synaptic current amplitude of approximately 1.

The synaptic current *I* of the *i*−th neuron is calculated as follows:

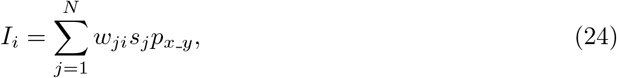

where *j* represents the index of a presynaptic neuron. *w*_*ji*_ is the weight of the synaptic connection from the *j*-th neuron to the *i*-th neuron. *N* is the total number of neurons. *x* and *y* indicate the classes of presynaptic and postsynaptic neurons, respectively, and the parameter *p*_*x−y*_ scales the synaptic current. As the extent to which the single spike of each class of neurons affects the membrane potential of different classes of postsynaptic neurons is not known clearly, the values of *p*_*x−y*_ were manually fitted such that the simulation results reproduce the experimental results as closely as possible. Here, the four LN subclasses share the same *p*_*x− y*_ value. First, the values of *p*_*x− y*_ where *y* is a PN or LN were set to reproduce the characteristics of oscillations observed in the antennal lobe *in vivo*. Next, the values of *p*_*x− y*_, where *y* is a KC, APL, or MBON-*α*1, were determined to make the responses of MBON-*α*1 as consistent as possible with the *in vivo* data. Finally, the values of *p*_*x− y*_, where *y* is MBON-*α*3 or SMP354 neuron, are set such that the success rate of olfactory associative learning becomes as high as possible. Note that, *p*_*x− y*_ is positive or negative when *x* is an excitatory or inhibitory neuron. All the parameter sets of neurons and synapses are listed in Supplementary Note 4.

### 4.6 FPGA implementation of the PQN model

In the FPGA implementation, the PQN model is simulated by the PQN engine. As an example, the details of the PQN engine of the PN mode are shown in Figure 9. Figure 9a shows the information flow of the PQN engine. The PQN engine updates the internal state variables and synaptic currents of 121 individual PNs in turn at each time step. The internal variables(*v, n, v*_*s*_, *u*, and *F*), input currents *I*, and synaptic currents *s* are sent via the PQN controller from block RAMs named PQN internal variables, I5, and *s*5 shown in Figure 6, respectively. The next step values of the internal variables and synaptic currents computed by the PQN engine are returned to the block RAMs and stored. Figure 9b shows a block diagram of part of the PQN engine of the PN mode. The full version is shown in Supplementary Figure 8. This circuit represents the calculation of the state variables *v* and *s*. The symbols×, +, and *M* in the figure represent the multipliers, adders, and multiplexers, respectively. Each state variable is computed in four pipelined stages. In the first stage, the square of *v* and the product of *u* and *I* are calculated using two multipliers. The second stage involves multiplication of the variables and coefficients determined from the parameters, and *v_x, s_S*, and *s_L* represent the results of the calculations, where *x* is *vv_S, vv_L, v_S, v_L, n, v*_*s*_, or *I*. For example, the calculation of *v_vv_S* is performed by multiplying the square of *v* by 0.021484375, the binary representation of which is 0.000001011. Therefore, the calculation of the sum of the sixth, eighth, and ninth right-shift operations on the square of *v* is performed (Fig. 9c). In the third stage, the results of the calculation of *v* and *s* are obtained by addition and selection based on the value of *v*. In the fourth stage, the value of *s* is determined based on the new value of *v*. When the old value of *v* is negative and the new value is zero or greater, the spike detector detects a spike. The internal variable *F* is not shown here since it is not used to update *v* and *s*, but is shown in the full version (see Supplementary Figure 8). The values of *v, n*, and *v*_*s*_ are updated every 1 ms, whereas the value of *u* is updated only once per second. The current firing frequency is calculated from the number of spikes counted in one second. The calculations for *n* and *v*_*s*_ are conducted in a manner similar to those for *v*. All state variables are expressed in an 18-bit fixed-point representation, of which 10 bits are the decimal part and the remaining are the integer part.

**Figure 9:**
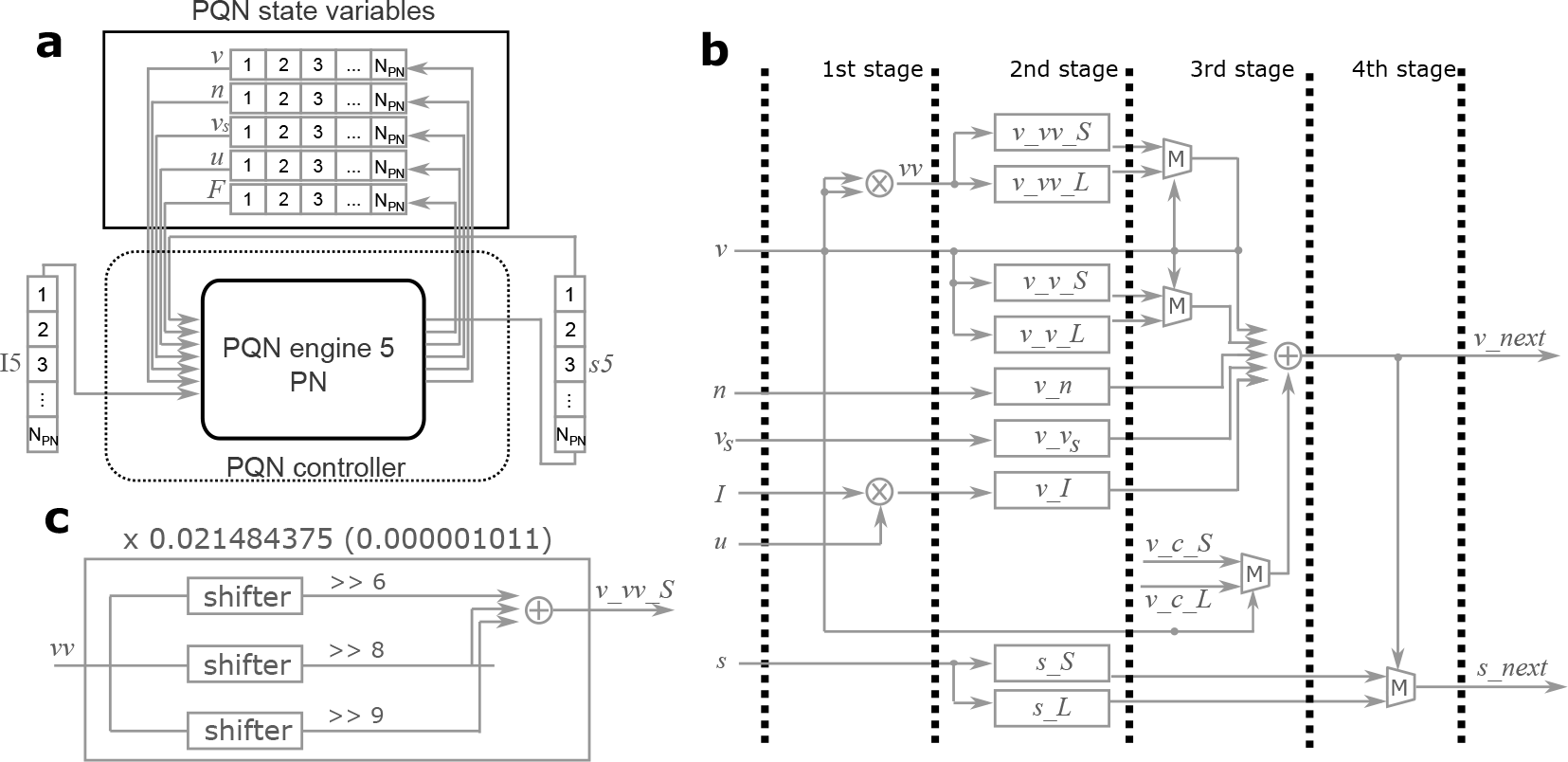
Details of the PQN engine of the PN mode. **a** Information flow of the PQN engine. N_PN_ is the number of PNs. **b** Block diagram of a part of the PQN engine. This circuit calculates the succeeding values of *v* and *s*. The symbols ×, +, and *M* represent multipliers, adders, and multiplexers, respectively. **c** Internal circuit for the calculation of *v_vv_S*.

## 5 Code availability

Codes and data are deposited in GitHub https://github.com/tnanami/fly-olfactory-network-fpga/tree/main.

## 6 Acknowledgements

This study was supported by the JSPS KAKENHI Grant Number 21K17849. H.K. was supported by a grant from RIKEN. T.H was supported by grants from National Institutes of Health (R01DC018874), National Science Foundation (2034783) and United States-Israel Binational Science Foundation (2019026). D.Y was supported by Toyobo Biotechnology Foundation Postdoctoral Fellowship and Japan Society for the Promotion of Science Overseas Research Fellowship. We thank Yoichi Seki and Kengo Inada for the recording data from LNs and KC.

## 7 Author information

### 7.1 Contributions

T.N, T.H, H.K and T.K conceptualized the work. T.N and T.K designed and developed the methodology. D.Y and M.S performed in vivo experiments. T.N performed in silico experiments and analyzed the data. T.N, T.H, H.K, and T.K wrote the paper.

### 7.2 Corresponding authors

Correspondence to Takuya Nanami

## References

[1] M. Kawato, “Internal models for motor control and trajectory planning.” Curr Opin Neurobiol, vol. 9, pp. 718–27, 01 1999.

[2] M. Frank, B. Loughry, and R. O’Reilly, “Frank mj, loughry b, o’reilly rc. interactions between frontal cortex and basal ganglia in working memory: a computational model. cogn affect behav neurosci 1: 137-160,” Cognitive, affective behavioral neuroscience, vol. 1, pp. 137–60, 07 2001.

[3] K. Norman and R. O’Reilly, “Modeling hippocampal and neocortical contributions to recognition memory: A complementary learning systems approach,” Psychological review, vol. 110, pp. 611–46, 11 2003.

[4] D. Walther and C. Koch, “Modeling attention to salient proto-objects,” NeuralNetworks, vol. 19, no. 9, pp. 1395–1407, 2006, brain and Attention. [Online]. Available: https://www.sciencedirect.com/science/article/pii/S0893608006002152

[5] P. Merolla, J. Arthur, R. Alvarez-Icaza, A. Cassidy, J. Sawada, F. Akopyan, B. Jackson, N. Imam, C. Guo, Y. Nakamura, B. Brezzo, I. Vo, S. Esser, R. Appuswamy, B. Taba, A. Amir, M. Flickner, W. Risk, R. Manohar, and D. Modh, “A million spiking-neuron integrated circuit with a scalable communication network and interface,” Science, vol. 345, no. 6197, pp. 668–673, August 2014.

[6] N. Qiao, H. Mostafa, F. Corradi, M. Osswald, F. Stefanini, D. Sumislawska, and G. Indiveri, “A reconfigurable on-line learning spiking neuromorphic processor comprising 256 neurons and 128k synapses,” Frontiers in Neuroscience, vol. 9, p. 141, 2015.

[7] M. Davies, N. Srinivasa, T. Lin, G. Chinya, Y. Cao, S. H. Choday, G. Dimou, P. Joshi, N. Imam, S. Jain, Y. Liao, C. Lin, A. Lines, R. Liu, D. Mathaikutty, S. McCoy, A. Paul, J. Tse, G. Venkataramanan, Y. Weng, A. Wild, Y. Yang, and H. Wang, “Loihi: A neuromorphic manycore processor with on-chip learning,” IEEE Micro, vol. 38, no. 1, pp. 82–99, 2018.

[8] H. Markram, E. Muller, S. Ramaswamy, M. Reimann, M. Abdellah, C. Aguado, A. Ailamaki, L. Alonso-Nanclares, N. Antille, S. Arsever, A. K. Guy Antoine, T. Berger, A. Bilgili, N. Buncic, A. Chalimourda, G. Chindemi, J.-D. Courcol, F. Delalondre, V. Delattre, and F. Schürmann, “Reconstruction and simulation of neocortical microcircuitry,” Cell, vol. 163, pp. 456–492, 10 2015.

[9] M. J. Bezaire, I. Raikov, K. Burk, D. Vyas, and I. Soltesz, “Interneuronal mechanisms of hippocampal theta oscillations in a full-scale model of the rodent ca1 circuit,” eLife, vol. 5, p. e18566, ec 2016.

[10] A. Ecker, A. Romani, S. Saray, S. Kali, M. Migliore, J. Falck, S. Lange, A. Mercer, A. M. Thomson, E. Muller, M. W. Reimann, and S. Ramaswamy, “Data-driven integration of hippocampal ca1 synaptic physiology in silico,” Hippocampus, vol. 30, no. 11, pp. 1129–1145, 2020.

[11] T. Nanami and T. Kohno, “Simple cortical and thalamic neuron models for digital arithmetic circuit implementation,” Frontiers in Neuroscience, section Neuromorphic Engineering, vol. 10, no. 181, 2016.

[12] T. Nanami and T. Kohno, “An fpga-based cortical and thalamic silicon neuronal network,” Journal of Robotics Networking and Artificial Life, vol. 2, no. 4, pp. 238–242, 2016.

[13] T. Nanami, K. Aihara, and T. Kohno, “Elliptic and parabolic bursting in a digital silicon neuron model,” in 2016 International Symposium on Nonlinear Theory and Its Applications, November 2016, pp. 198–201.

[14] T. Nanami, F. Grassia, and T. Kohno, “A parameter optimization method for digital spiking silicon neuron model,” Journal of Robotics Networking and Artificial Life, vol. 4, no. 1, pp. 97–101, 2017.

[15] T. Nanami, F. Grassia, and T. Kohno, “A metaheuristic approach for parameter fitting in digital spiking silicon neuron model,” Journal of Robotics Networking and Artificial Life, vol. 5, no. 1, pp. 32–36, 2018.

[16] T. Nanami and T. Kohno, “Piecewise quadratic neuron model: A tool for close-to-biology spiking neuronal network simulation on dedicated hardware,” Frontiers in Neuroscience, vol. 16, 2023. [Online]. Available: https://www.frontiersin.org/articles/10.3389/fnins.2022.1069133

[17] E. M. Izhikevich, “Simple model of spiking neurons,” IEEE Trans. Neural Networks, pp. 1569–1572, 2003.

[18] R. Brette and W. Gerstner, “Adaptive Exponential Integrate-and-Fire Model as an Effective Description of Neuronal Activity,” J. Neurophysiol., vol. 94, pp. 3637–3642, 2005.

[19] H. Alle and J. R. P. Geiger, “Combined analog and action potential coding in hippocampal mossy fibers,” Science, vol. 311, pp. 1290–1293, 2006.

[20] A. L. Hodgkin, “The local electric changes associated with repetitive action in a non-medullated axon.” The Journal of physiology, vol. 107, no. 2, pp. 165–181, Mar. 1948.

[21] L. K. Scheffer, C. S. Xu, M. Januszewski, Z. Lu, S.-y. Takemura, K. J. Hayworth, G. B. Huang, K. Shinomiya, J. Maitlin-Shepard, S. Berg, J. Clements, P. M. Hubbard, W. T. Katz, L. Umayam, T. Zhao, D. Ackerman, T. Blakely, J. Bogovic, T. Dolafi, D. Kainmueller, T. Kawase, K. A. Khairy, L. Leavitt, P. H. Li, L. Lindsey, N. Neubarth, D. J. Olbris, H. Otsuna, E. T. Trautman, M. Ito, A. S. Bates, J. Goldammer, T. Wolff, R. Svirskas, P. Schlegel, E. Neace, C. J. Knecht, C. X. Alvarado, D. A. Bailey, S. Ballinger, J. A. Borycz, B. S. Canino, N. Cheatham, M. Cook, M. Dreher, O. Duclos, B. Eubanks, K. Fairbanks, S. Finley, N. Forknall, A. Francis, G. P. Hopkins, E. M. Joyce, S. Kim, N. A. Kirk, J. Kovalyak, S. A. Lauchie, A. Lohff, C. Maldonado, E. A. Manley, S. McLin, C. Mooney, M. Ndama, O. Ogundeyi, N. Okeoma, C. Ordish, N. Padilla, C. M. Patrick, T. Paterson, E. E. Phillips, E. M. Phillips, N. Rampally, C. Ribeiro, M. K. Robertson, J. T. Rymer, S. M. Ryan, M. Sammons, A. K. Scott, A. L. Scott, A. Shinomiya, C. Smith, K. Smith, N. L. Smith, M. A. Sobeski, A. Suleiman, J. Swift, S. Takemura, I. Talebi, D. Tarnogorska, E. Tenshaw, T. Tokhi, J. J. Walsh, T. Yang, J. A. Horne, F. Li, R. Parekh, P. K. Rivlin, V. Jayaraman, M. Costa, G. S. Jefferis, K. Ito, S. Saalfeld, R. George, I. A. Meinertzhagen, G. M. Rubin, H. F. Hess, V. Jain, and S. M. Plaza, “A connectome and analysis of the adult Drosophila central brain,” eLife, vol. 9, p. e57443, sep 2020. [Online]. Available: 10.7554/eLife.57443

[22] R. I. Wilson, “Early olfactory processing in drosophila: Mechanisms and principles,” Annual Review of Neuroscience, vol. 36, no. 1, pp. 217–241, 2013.

[23] M. Modi, Y. Shuai, and G. Turner, “The drosophila mushroom body: From architecture to algorithm in a learning circuit,” Annual Review of Neuroscience, vol. 43, 07 2020.

[24] J. Wessnitzer, J. M. Young, J. D. Armstrong, and B. Webb, “A model of non-elemental olfactory learning in drosophila,” Journal of Computational Neuroscience, vol. 32, pp. 197– 212, 04 2012.

[25] F. Faghihi, A. A. Moustafa, R. Heinrich, and F. Worgotter, “A computational model of conditioning inspired by drosophila olfactory system,” Neural Networks, vol. 87, pp. 96–108, 2017.

[26] A. Kennedy, “Learning with naturalistic odor representations in a dynamic model of the drosophila olfactory system,” bioRxiv, 2019.

[27] N. K. Tanaka, K. Ito, and M. Stopfer, “Odor-evoked neural oscillations in drosophila are mediated by widely branching interneurons,” Journal of Neuroscience, vol. 29, no. 26, pp. 8595–8603, 2009.

[28] G. C. Turner, M. Bazhenov, and G. Laurent, “Olfactory representations by drosophila mushroom body neurons,” Journal of Neurophysiology, vol. 99, no. 2, pp. 734–746, 2008.

[29] T. Hige, Y. Aso, M. N. Modi, G. M. Rubin, and G. Turner, “Heterosynaptic plasticity underlies aversive olfactory learning in drosophila,” Neuron, vol. 88, pp. 985–998, 12 2015.

[30] M. Bazhenov, M. Stopfer, M. Rabinovich, R. Huerta, H. D. Abarbanel, T. J. Sejnowski, and G. Laurent, “Model of transient oscillatory synchronization in the locust antennal lobe,” Neuron, vol. 30, no. 2, pp. 553–567, 2001.

[31] M. Bazhenov, M. Stopfer, M. Rabinovich, H. Abarbanel, T. Sejnowski, and G. Laurent, “Model of cellular and network mechanisms for odor-evoked temporal patterning in the locust antennal lobe,” Neuron, vol. 30, pp. 569–81, 06 2001.

[32] Y. Seki, J. Rybak, D. Wicher, S. Sachse, and B. S. Hansson, “Physiological and morphological characterization of local interneurons in the drosophila antennal lobe,” Journal of Neurophysiology, vol. 104, no. 2, pp. 1007–1019, 2010.

[33] K. Inada, Y. Tsuchimoto, and H. Kazama, “Origins of cell-type-specific olfactory processing in the drosophila mushroom body circuit,” Neuron, vol. 95, no. 2, pp. 357–367.e4, 2017.

[34] R. Stocker, M. Lienhard, A. Borst, and K. Fischbach, “Neuronal architecture of the antennal lobe in drosophila melanogaster,” Cell and Tissue Research, vol. 262, pp. 9–34, 1990.

[35] Y. Aso, D. Hattori, Y. Yu, R. M. Johnston, N. A. Iyer, T.-T. Ngo, H. Dionne, L. Abbott, R. Axel, H. Tanimoto, and G. M. Rubin, “The neuronal architecture of the mushroom body provides a logic for associative learning,” eLife, vol. 3, 2014.

[36] Y. Aso, D. Yamada, D. Bushey, K. Hibbard, M. Sammons, H. Otsuna, Y. Shuai, and T. Hige, “Neural circuit mechanisms for transforming learned olfactory valences into wind-oriented movement,” bioRxiv, 2022. [Online]. Available: https://www.biorxiv.org/content/early/2022/12/24/2022.12.21.521497

[37] “neuPrint, hemibrain: v1.0.1,” https://neuprint.janelia.org/.

[38] Y.-H. Chou, M. Spletter, E. Yaksi, J. Leong, R. Wilson, and L. Luo, “Diversity and wiring variability of olfactory local interneurons in the drosophila antennal lobe,” Natureneuroscience, vol. 13, pp. 439–49, 02 2010.

[39] R. Storn and K. Price, “A simple and efficient heuristic for global optimization over continuous spaces,” Journal of Global Optimization, vol. 11, no. 4, pp. 341–359, 1997.

[40] R. I. Wilson, G. C. Turner, and G. Laurent, “Transformation of olfactory representations in the drosophila antennal lobe,” Science, vol. 303, no. 5656, pp. 366–370, 2004.

[41] M. Stopfer, S. Bhagavan, B. Smith, and G. Laurent, “Impaired odour discrimination on desynchronization of odour-encoding neural assemblies,” Nature, vol. 390, pp. 70–4, 12 1997.

[42] J. Perez-Orive, O. Mazor, G. C. Turner, S. Cassenaer, R. I. Wilson, and G. Laurent, “Oscillations and sparsening of odor representations in the mushroom body,” Science, vol. 297, no. 5580, pp. 359–365, 2002.

[43] M. Tsodyks and H. Markram, “The neural code between neocortical pyramidal neurons depends on neurotransmitter release probability,” Proceedings of the National Academy of Sciences, vol. 94, no. 2, pp. 719–723, 1997.

[44] M. Heisenberg, “Heisenberg, m. mushroom body memoir: from maps to models. nat. rev. neurosci. 4, 266-275,” Nature reviews. Neuroscience, vol. 4, pp. 266–75, 05 2003.

[45] K. Akbarzadeh-Sherbaf, B. Abdoli, S. Safari, and A.-H. Vahabie, “A scalable fpga architecture for randomly connected networks of hodgkin-huxley neurons,” Frontiers in Neuroscience, vol. 12, p. 698, 2018.

[46] F. Khoyratee, F. Grassia, S. Saighi, and T. Levi, “Optimized real-time biomimetic neural network on fpga for bio-hybridization,” Frontiers in Neuroscience, vol. 13, p. 377, 2019.

[47] A. Amara, F. Amiel, and T. Ea, “Fpga vs. asic for low power applications,” Microelectronics Journal, vol. 37, no. 8, pp. 669–677, 2006.

[48] I. Kuon and J. Rose, “Measuring the gap between fpgas and asics,” IEEE Transactions on Computer-Aided Design of Integrated Circuits and Systems, vol. 26, no. 2, pp. 203–215, 2007.

[49] D. Münch and C. Galizia, “Door 2.0 - comprehensive mapping of drosophila melanogaster odorant responses,” Scientific Report, vol. 6, p. 21841, 2016.

[50] H. Kazama and R. Wilson, “Origins of correlated activity in an olfactory circuit,” Natureneuroscience, vol. 12, pp. 1136–44, 09 2009.

[51] V. Bhandawat, S. Olsen, N. Gouwens, M. Schlief, and R. Wilson, “Sensory processing in the drosophila antennal lobe increases reliability and separability of ensemble odor representations,” Nature neuroscience, vol. 10, pp. 1474–82, 12 2007.

[52] S. Olsen and R. Wilson, “Lateral presynaptic inhibition mediates gain control in an olfactory circuit,” Nature, vol. 452, pp. 956–60, 05 2008.

[53] S. Caron, V. Ruta, L. Abbott, and R. Axel, “Random convergence of olfactory inputs in the drosophila mushroom body,” Nature, vol. 497, 04 2013.

[54] F. Li, J. W. Lindsey, E. C. Marin, N. Otto, M. Dreher, G. Dempsey, I. Stark, A. S. Bates, M. W. Pleijzier, P. Schlegel, A. Nern, S.-y. Takemura, N. Eckstein, T. Yang, A. Francis, A. Braun, R. Parekh, M. Costa, L. K. Scheffer, Y. Aso, G. S. Jefferis, L. F. Abbott, A. Litwin-Kumar, S. Waddell, and G. M. Rubin, “The connectome of the adult drosophila mushroom body provides insights into function,” eLife, vol. 9, p. e62576, ec 2020. [Online]. Available: 10.7554/eLife.62576

[55] Y. Aso, D. Sitaraman, T. Ichinose, K. R. Kaun, K. Vogt, G. Belliart-Guerin, P.-Y. Placais, A. A. Robie, N. Yamagata, C. Schnaitmann, W. J. Rowell, R. M. Johnston, T.-T. B. Ngo, N. Chen, W. Korff, M. N. Nitabach, U. Heberlein, T. Preat, K. M. Branson, H. Tanimoto, and G. M. Rubin, “Mushroom body output neurons encode valence and guide memory-based action selection in Drosophila,” eLife, vol. 3, 2014.

[56] A. Matheson, A. Lanz, A. Licata, T. Currier, M. Syed, and K. Nagel, “A neural circuit for wind-guided olfactory navigation,” Nature communications, 04 2021.

[57] Y. Aso and G. M. Rubin, “Dopaminergic neurons write and update memories with cell-type-specific rules,” eLife, vol. 5, p. e16135, jul 2016. [Online]. Available: 10.7554/eLife.16135

[58] T. Tully and W. Quinn, “Classical-conditioning and retention in normal and mutant drosophila-melanogaster,” Journal of comparative physiology. A, Sensory, neural, and behavioral physiology, vol. 157, pp. 263–77, 10 1985.

[59] R. Cohn, I. Morantte, and V. Ruta, “Coordinated and compartmentalized neuromodulation shapes sensory processing in drosophila,” Cell, vol. 163, pp. 1742–1755, 12 2015.

[60] D. Owald, J. Felsenberg, C. Talbot, G. Das, E. Perisse, W. Huetteroth, and S. Waddell, “Activity of defined mushroom body output neurons underlies learned olfactory behavior in drosophila,” Neuron, vol. 86, 04 2015.

[61] A. C. von Philipsborn, T. Liu, J. Y. Yu, C. Masser, S. S. Bidaye, and B. J. Dickson, “Neuronal control of drosophila courtship song,” Neuron, vol. 69, no. 3, pp. 509–522, 2011. [Online]. Available: https://www.sciencedirect.com/science/article/pii/S0896627311000572

[62] H. Inagaki, Y. Jung, E. Hoopfer, A. Wong, N. Mishra, J. Lin, R. Tsien, and D. Anderson, “Optogenetic control of drosophila using a red-shifted channelrhodopsin reveals experiencedependent influences on courtship,” Nature methods, vol. 11, 12 2013.

[63] M. de Bruyne, P. J. Clyne, and J. R. Carlson, “Odor coding in a model olfactory organ: Thedrosophila maxillary palp,” Journal of Neuroscience, vol. 19, no. 11, pp. 4520–4532, 1999.

[64] E. A. Hallem and J. R. Carlson, “Coding of odors by a receptor repertoire,” Cell, vol. 125, no. 1, pp. 143–160, 2006.

[65] H. Kazama and R. I. Wilson, “Homeostatic matching and nonlinear amplification at identified central synapses,” Neuron, vol. 58, no. 3, pp. 401–413, 2008. [Online]. Available: https://www.sciencedirect.com/science/article/pii/S0896627308001840

[66] R. FitzHugh, “Impulses and physiological states in theoretical models of nerve membrane,” j-BIOPHYS-J, vol. 1, pp. 445–466, 1961.

[67] J. S. Nagumo, S. Arimoto, and S. Yoshizawa, “An active pulse transmission line simulating nerve axon,” j-PROC-IRE, vol. 50, pp. 2061–2071, 1962.

[68] J. L. Hindmarsh and R. M. Rose, “A Model of Neuronal Bursting Using Tree Coupled First Order Differential Equations,” Philos Trans Royal Soc London, vol. B221, pp. 87–102, 1984.

[69] N. W. Gouwens and R. I. Wilson, “Signal propagation in drosophila central neurons,” Journal of Neuroscience, vol. 29, no. 19, pp. 6239–6249, 2009. [Online]. Available: https://www.jneurosci.org/content/29/19/6239

[70] H. Amin, A. A. Apostolopoulou, R. Suarez-Grimalt, E. Vrontou, and A. C. Lin, “Localized inhibition in the Drosophila mushroom body,” Elife, vol. 9, p. e56954, sep 2020.

[71] G. G. Turrigiano, “Homeostatic plasticity in neuronal networks: the more things change, the more they stay the same,” Trends in Neurosciences, vol. 22, no. 5, pp. 221–227, 1999. [Online]. Available: https://www.sciencedirect.com/science/article/pii/S0166223698013411

[72] J. Li, Y. Katori, and T. Kohno, “An fpga-based silicon neuronal network with selectable excitability silicon neurons,” Frontiers in neuroscience, vol. 6, no. 183, 2012.

[73] D. Lee, H. Su, and D. K. O’Dowd, “Gaba receptors containing rdl subunits mediate fast inhibitory synaptic transmission in drosophila neurons,” Journal of Neuroscience, vol. 23, no. 11, pp. 4625–4634, 2003.

[74] H. Gu and D. K. O’Dowd, “Cholinergic synaptic transmission in adult drosophila kenyon cells in situ,” Journal of Neuroscience, vol. 26, no. 1, pp. 265–272, 2006.

