## Supplemental for "A lightweight data-driven spiking neural network model of *Drosophila* olfactory nervous system with dedicated hardware support"

**2 Supplementary Note 1** **10**
**3 Supplementary Note 2** **11**
**4 Supplementary Note 3** **11**
**5 Supplementary Note 4** **12**

### 1 Supplementary Figures

#### 1.1 Supplementary Figure 1. Responses of Krasavietz class1 in vivo and in silico

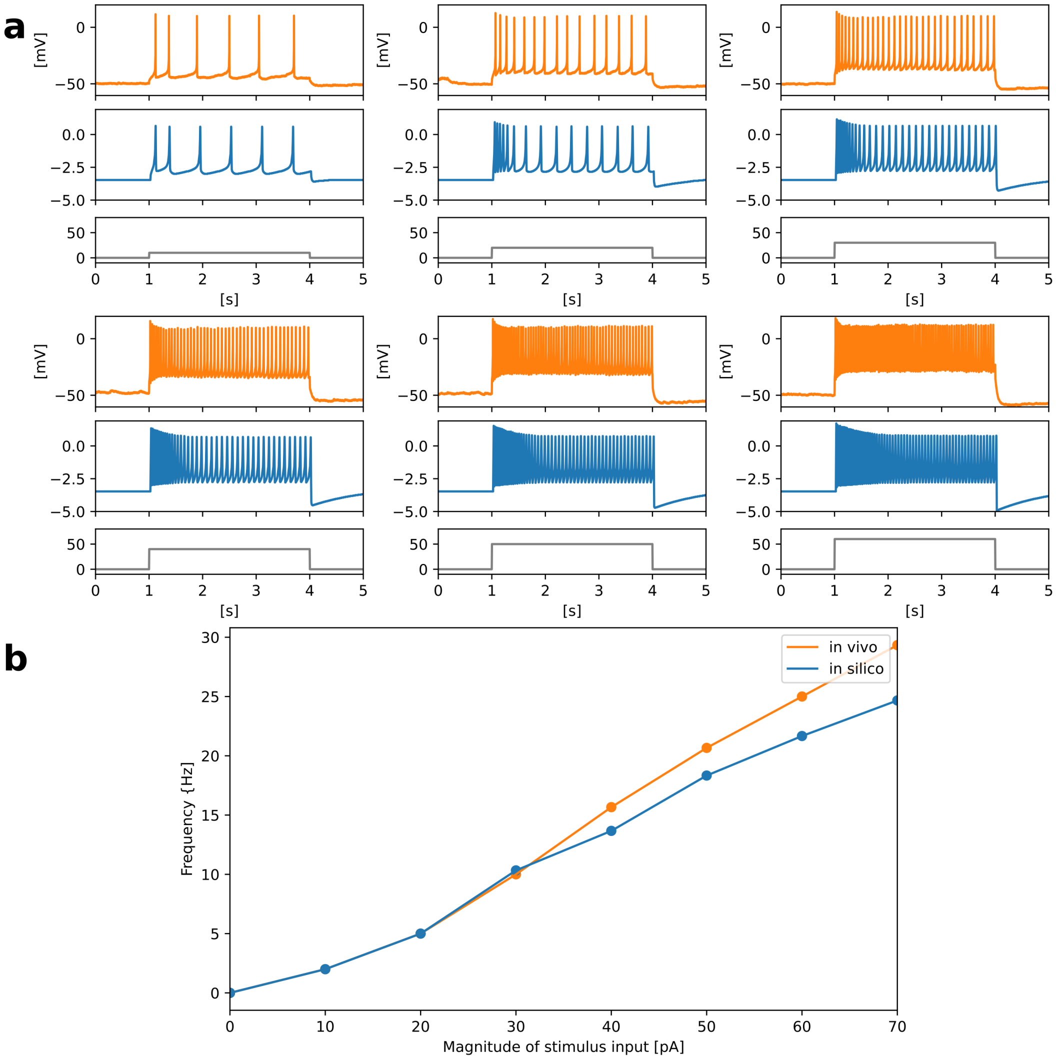

Supplementary Figure 1: Responses of Krasavietz class1 in vivo and in silico. **a** Responses of somatic membrane potentials in vivo (orange) and in silico (blue) in response to step stimulus inputs of several magnitudes. **b** Transition of firing frequency. The horizontal axis represents the magnitude of stimulus input, and the vertical axis represents the frequency.

#### 1.2 Supplementary Figure 2. Responses of Krasavietz class2 in vivo and in silico

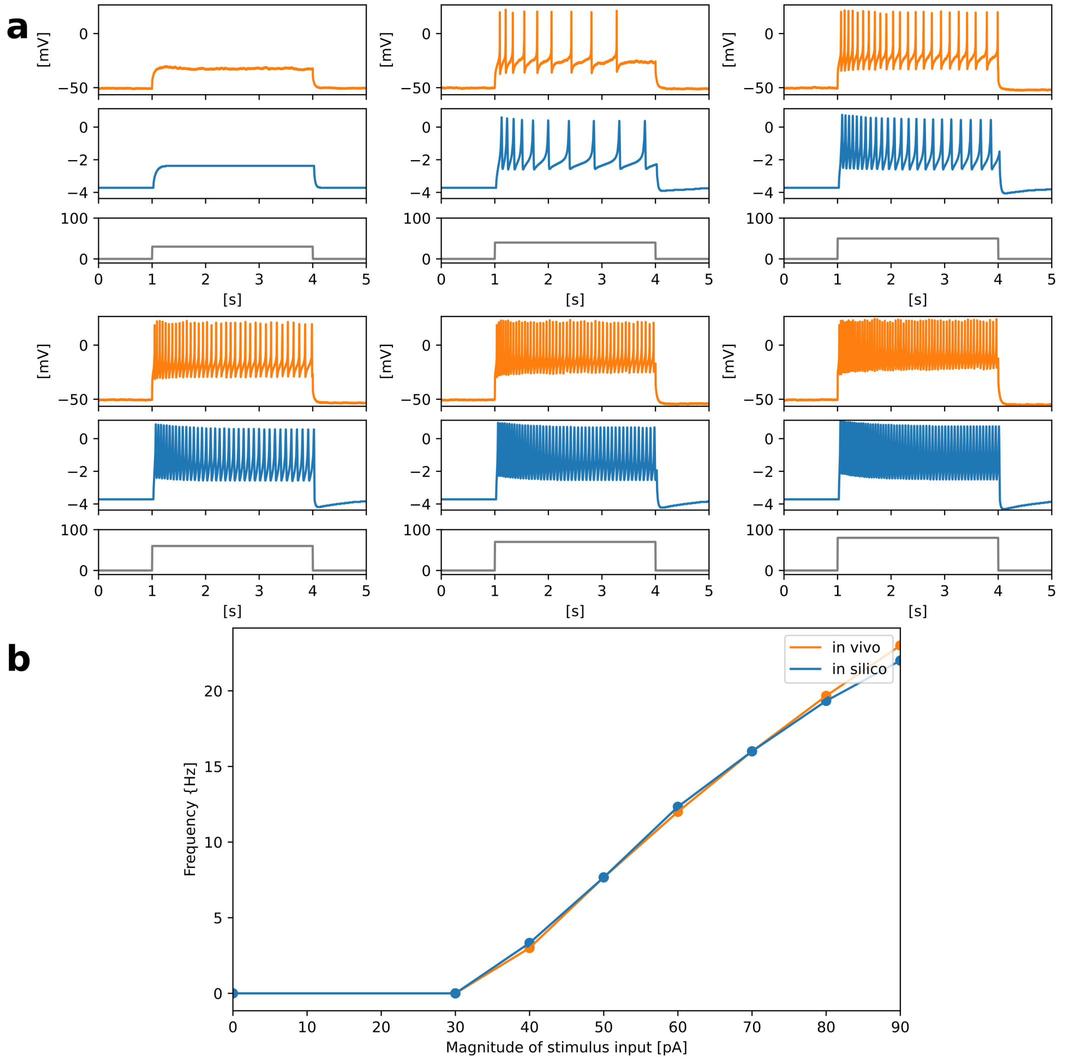

Supplementary Figure 2: Responses of Krasavietz class2 in vivo and in silico. **a** Responses of somatic membrane potentials in vivo (orange) and in silico (blue) in response to step stimulus inputs of several magnitudes. **b** Transition of firing frequency. The horizontal axis represents the magnitude of stimulus input, and the vertical axis represents the frequency.

#### 1.3 Supplementary Figure 3. Responses of NP1227 class1 in vivo and in silico

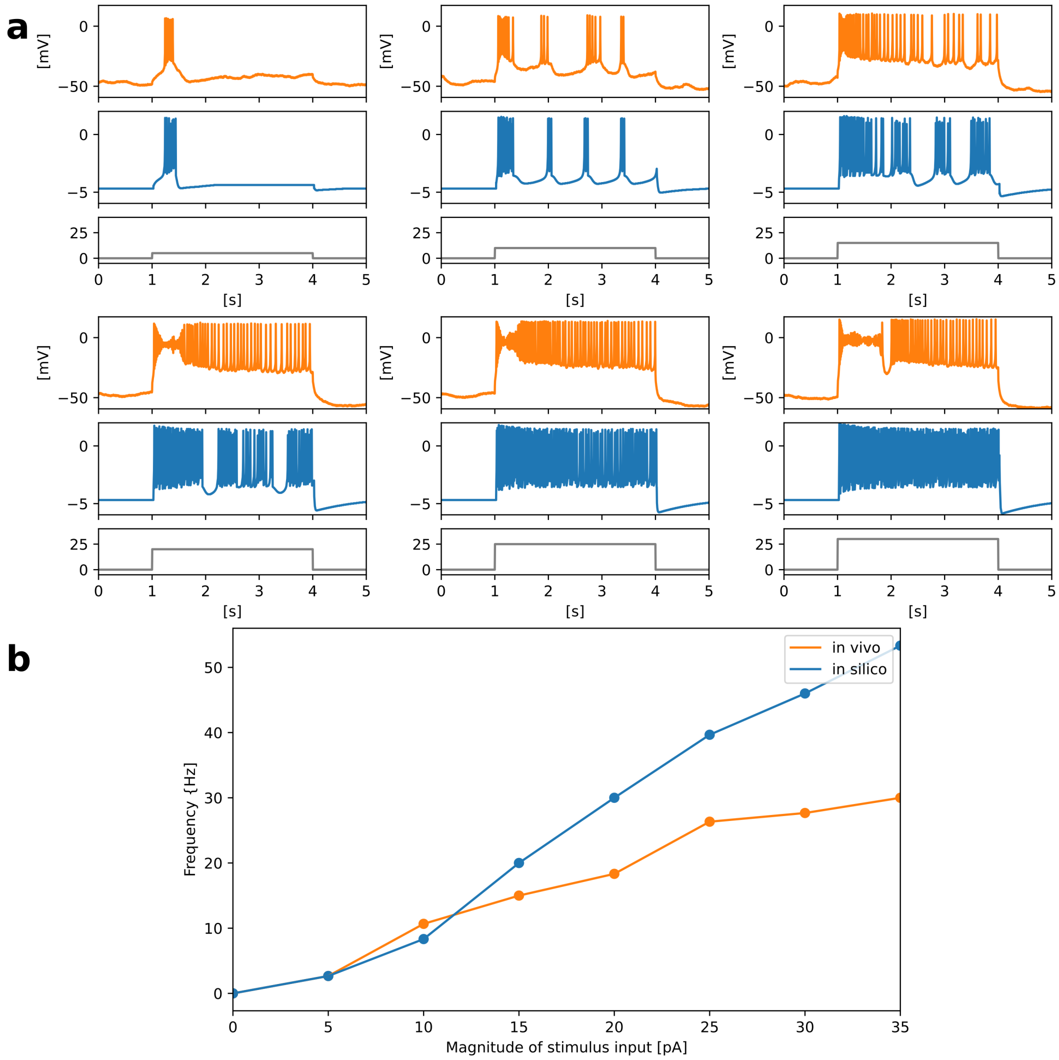

Supplementary Figure 3: Responses of NP1227 class1 in vivo and in silico. **a** Responses of somatic membrane potentials in vivo (orange) and in silico (blue) in response to step stimulus inputs of several magnitudes. **b** Transition of firing frequency. The horizontal axis represents the magnitude of stimulus input, and the vertical axis represents the frequency.

#### 1.4 Supplementary Figure 4. Responses of NP2426 class1 in vivo and in silico

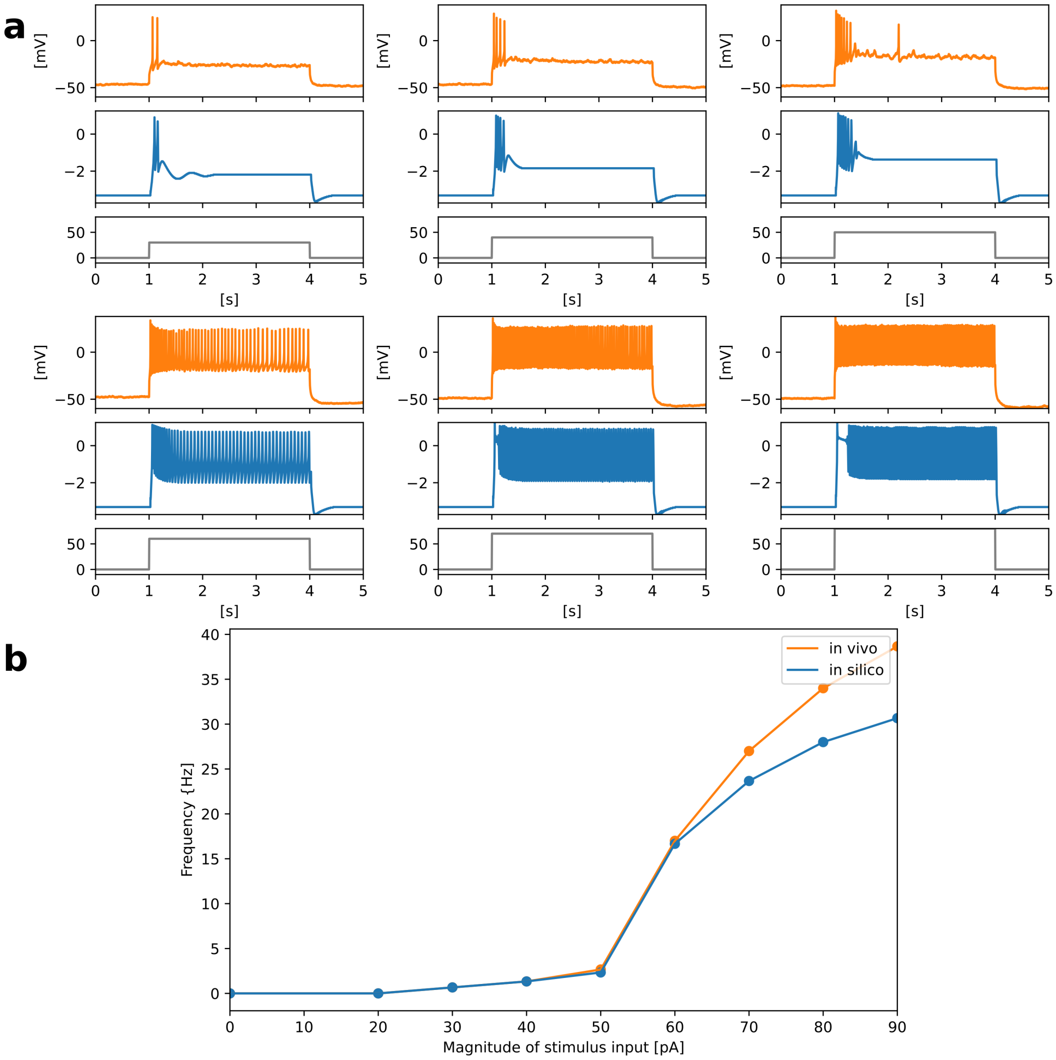

Supplementary Figure 4: Responses of NP2426 class1 in vivo and in silico. **a** Responses of somatic membrane potentials in vivo (orange) and in silico (blue) in response to step stimulus inputs of several magnitudes. **b** Transition of firing frequency. The horizontal axis represents the magnitude of stimulus input, and the vertical axis represents the frequency.

#### 1.5 Supplementary Figure 5. Responses of PN in vivo and in silico

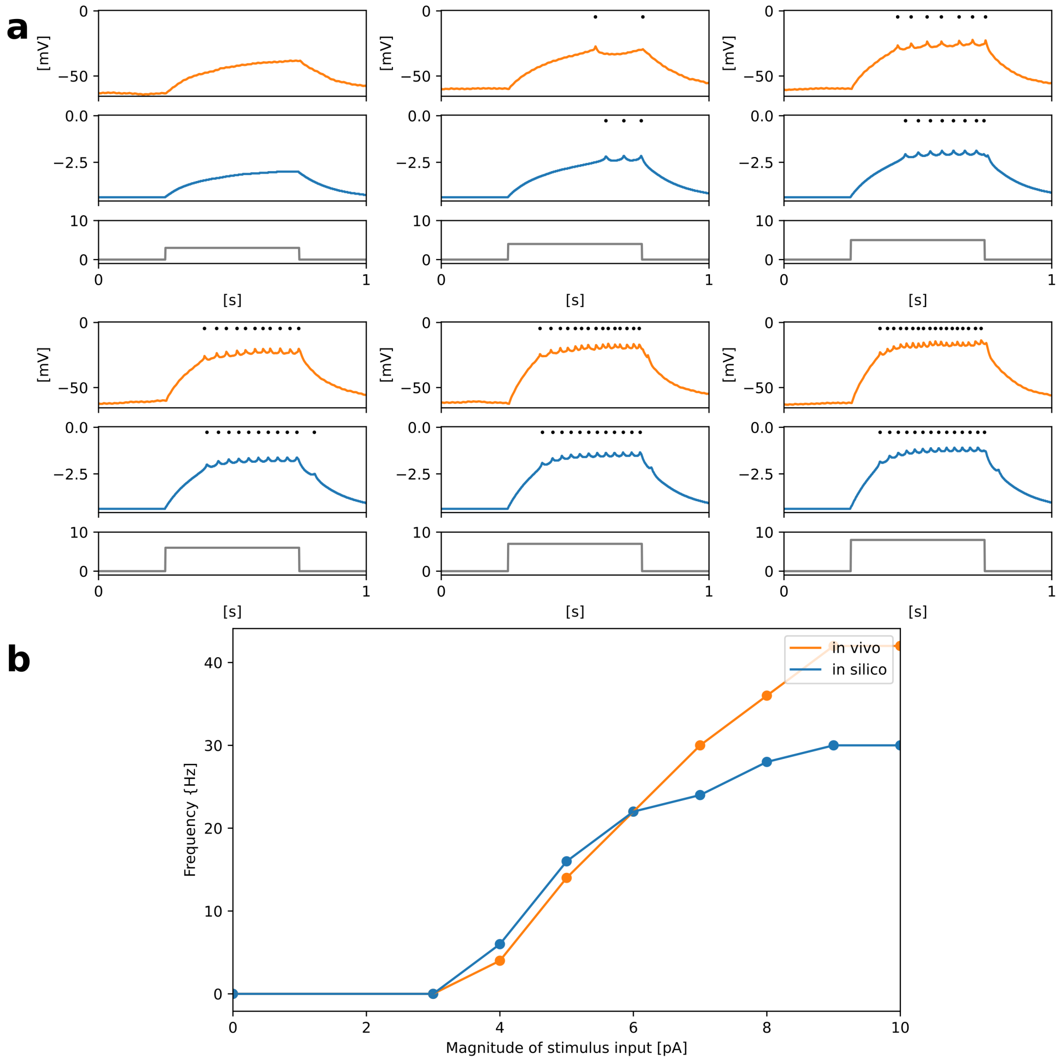

Supplementary Figure 5: Responses of PN in vivo and in silico. **a** Responses of somatic membrane potentials in vivo (orange) and in silico (blue) in response to step stimulus inputs of several magnitudes. **b** Transition of firing frequency. The horizontal axis represents the magnitude of stimulus input, and the vertical axis represents the frequency.

#### 1.6 Supplementary Figure 6. Responses of KC in vivo and in silico

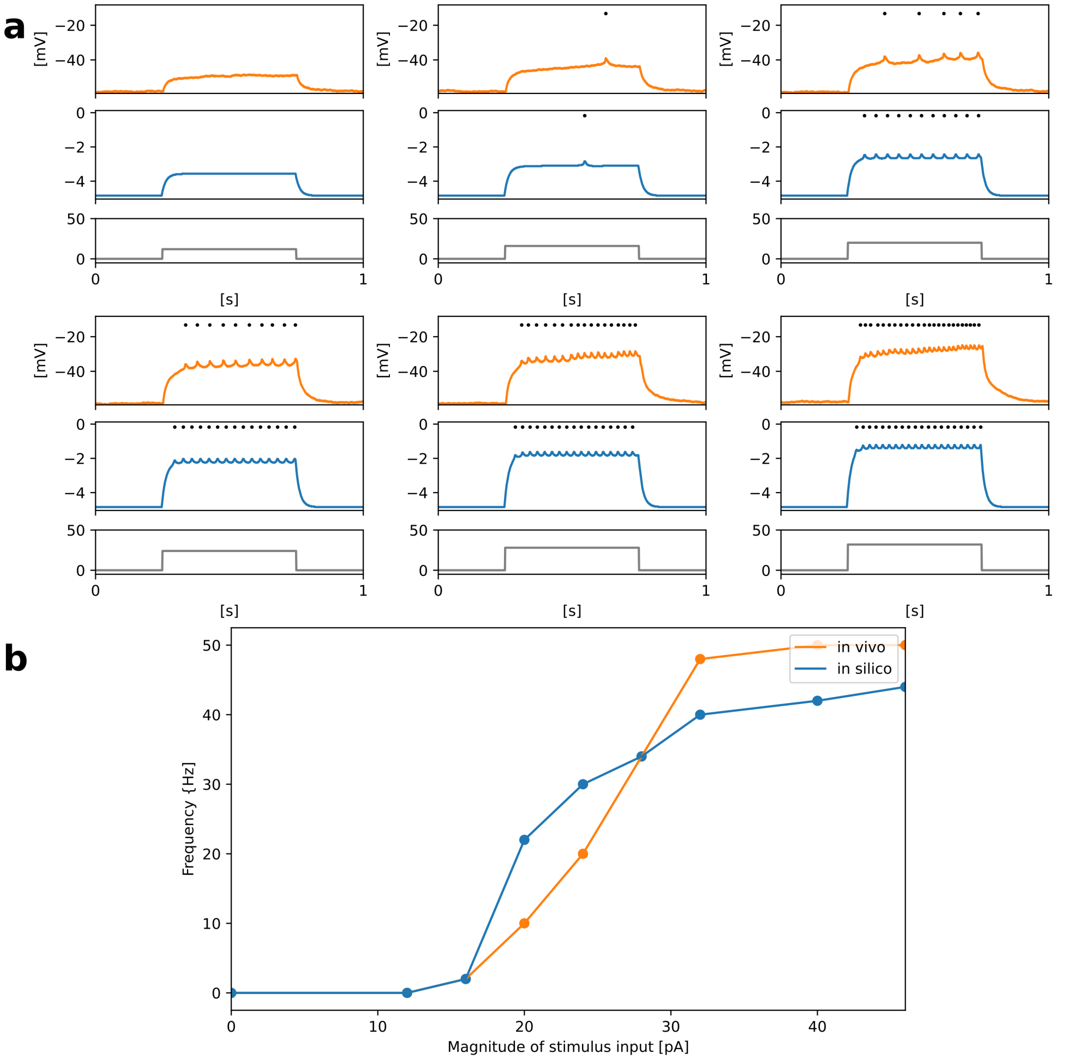

Supplementary Figure 6: Responses of KC in vivo and in silico. **a** Responses of somatic membrane potentials in vivo (orange) and in silico (blue) in response to step stimulus inputs of several magnitudes. **b** Transition of firing frequency. The horizontal axis represents the magnitude of stimulus input, and the vertical axis represents the frequency.

#### 1.7 Supplementary Figure 7. Responses of MBON in vivo and in silico

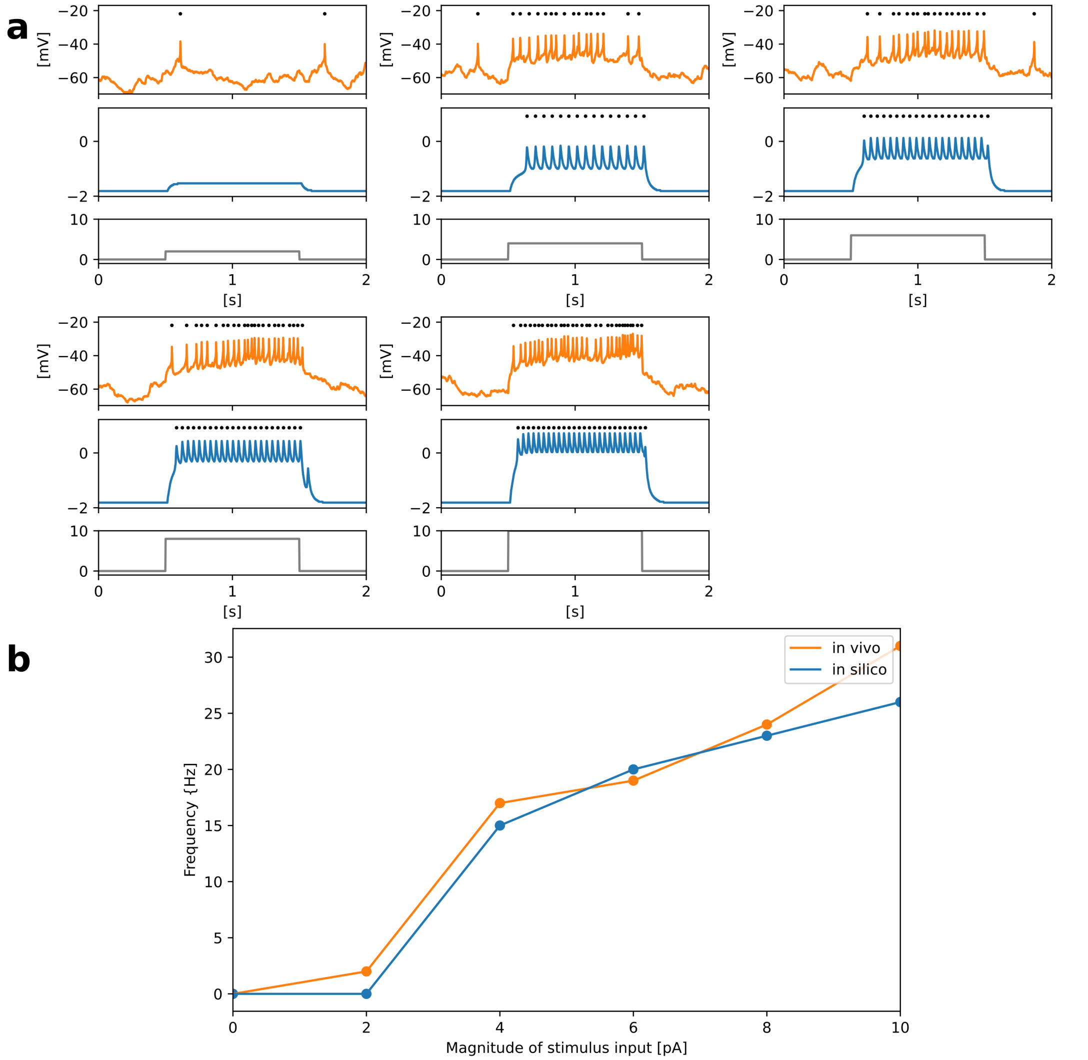

Supplementary Figure 7: Responses of MBON in vivo and in silico. **a** Responses of somatic membrane potentials in vivo (orange) and in silico (blue) in response to step stimulus inputs of several magnitudes. **b** Transition of firing frequency. The horizontal axis represents the magnitude of stimulus input, and the vertical axis represents the frequency.

#### 1.8 Supplementary Figure 8. Block diagram of the PQN engine of the PN mode

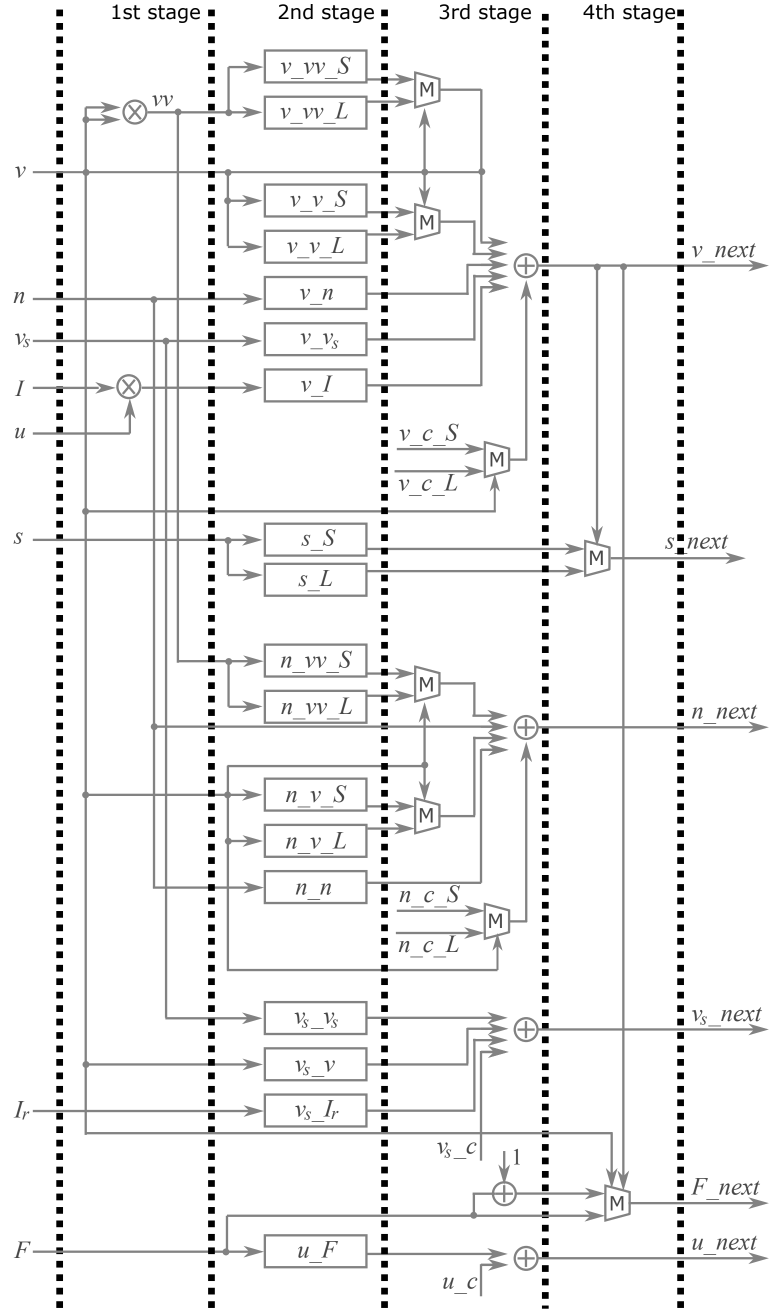

Supplementary Figure 8: A block diagram of the PQN engine of the PN mode.

##### 1.9 Supplementary Figure 9. Averages of the peak power spectra of PNs in response to six odors

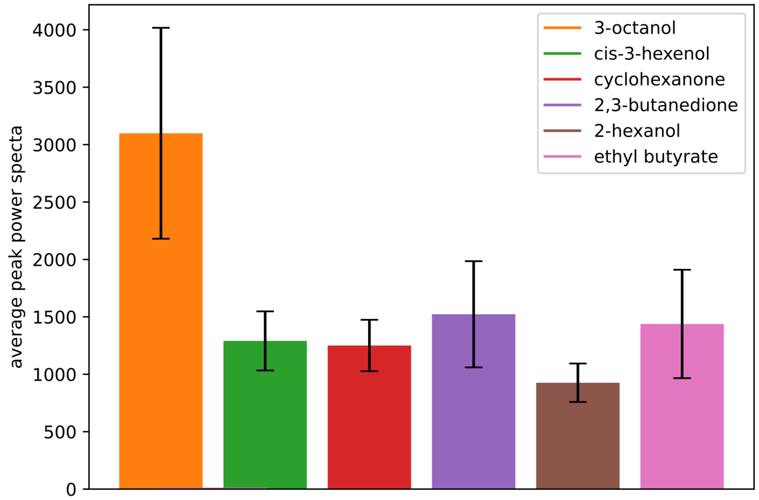

Supplementary Figure 9: Averages of the peak power spectra of PNs when one of the six odorants, 3-octanol, cis-3-hexenol, cyclohexanone, 2,3-butanedione, 2-hexanol, and ethyl butyrate, was applied. Error bars represent standard deviation over five trials.

#### 2 Supplementary Note 1

ORNs, PNs, KCs, and MBON-$\alpha3$ are cholinergic [1][2][3][4] and form excitatory synapses. Some LNs are cholinergic [5] or glutamatergic [6] and are considered as sources of excitatory or inhibitory input [5][7]. However, most LNs are GABAergic [8][9] and have been shown to provide inhibitory input [10][11]. Thus, in this model, all LNs were set as inhibitory. APL is GABAergic [3] and inhibitory, whereas MBON-$\alpha1$ is glutaminergic [12] and inhibitory.

### 3 Supplementary Note 2

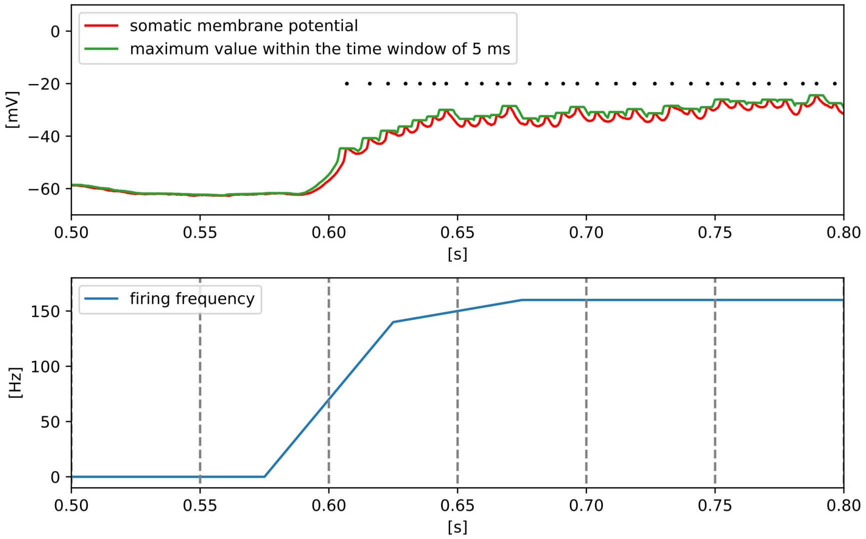

Supplementary Figure 10: Spike detection and frequency calculation for in vivo data of the somatic membrane potential.

Here, we describe the detection of spike timing from the somatic membrane potentials of in vivo data. The amplitude of spikes observed in the soma is decayed and small. In addition, the baseline of the somatic membrane potential during repeated firings fluctuates significantly. Therefore, we first plotted the maximum value of the membrane potential within the time window of 5 ms at each time point (Supplementary Fig. 10). Here, a spike is detected at a time point when the value of the time point is equal to the maximum value within the time window and the value of the time point is greater than the value of the previous time point. The latter rule prevents the detection of spikes twice when two adjacent maxima of the same value are measured at the top of the spike. Then, for each 50ms, the number of spikes was counted, and the instantaneous firing frequency was calculated as follows:

$$\text{firing frequency} = \mathrm{n_0/w_0} \tag{1}$$

where $n_0$ is the number of spikes in the 50 ms period and $w_0$ is 50 ms.

In the simulation, a spike was detected when the value of the membrane potential of the axonal compartment exceeded 0. The calculation of the frequency was performed in the same way as that in the in vivo data.

### 4 Supplementary Note 3

Before conducting all the experiments shown in Figures 3–5, 300 seconds of ORN input data were provided to the network to implement the homeostatic control of synaptic input observed in PNs. The data structure was identical to that used for olfactory associative learning, where six odorants, 3-octanol, cis-3-hexenol, cyclohexanone, 2,3-butanedione, 2-hexanol, and ethyl butyrate, were applied sequentially for one second every five seconds. Supplementary Figure

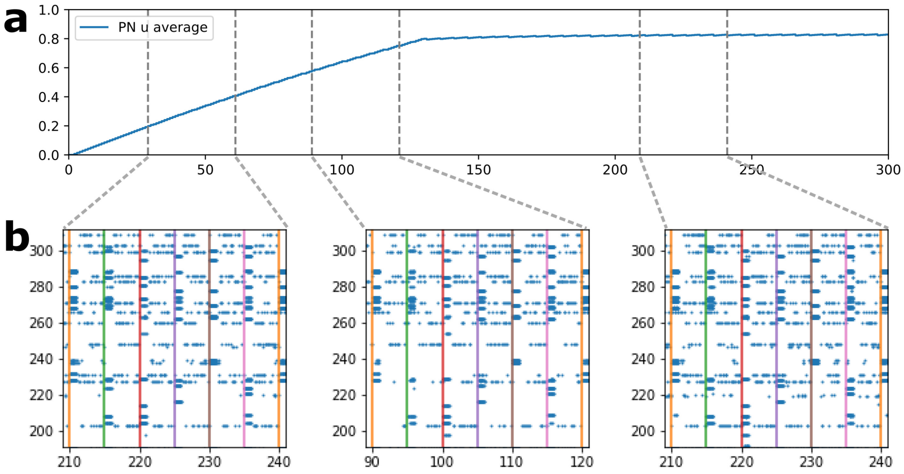

Supplementary Figure 11: Transition of the averaged value of $u$ for all PNs during the homeostatic period.

11a shows the transition of the average value of $u$ for all the PNs during this period. The initial value of $u$ is fixed at 0, and as the value gradually increases, a larger number of PNs fire to the odor (Supplementary Fig. 11b).

### 5 Supplementary Note 4

Table 1: Parameter set for the Krasavietz class1.

| Par. | Value | Par. | Value |
|---|---|---|---|
| $\Delta$ t | 0.001 | $\tau$ | 0.008 |
| $afn$ | 1.021484375 | $afp$ | -2.3544921875 |
| $bfn$ | -1.09375 | $cfn$ | 0.93359375 |
| $agn$ | -0.7119140625 | $agp$ | 9.5595703125 |
| $bgn$ | -4.8232421875 | $cgn$ | 13.427734375 |
| $ahn$ | -0.7041015625 | $ahp$ | 8.640625 |
| $bhn$ | -3.6533203125 | $chn$ | 4.521484375 |
| $I_{b0}$ | 5.2509765625 | $k_I$ | 16.3193359375 |
| $\phi$ | 0.703125 | $\epsilon_q$ | 0.0107421875 |
| $r_g$ | -1.234375 | $r_h$ | -1.955078125 |
| $m_0$ | -1 | $m_1$ | 1 |

### References

[1] K. Yasuyama and P. M. Salvaterra, “Localization of choline acetyltransferase-expressing neurons in drosophila nervous system,” Microscopy Research and Technique, vol. 45, no. 2, pp. 65–79, 1999.

Table 2: Parameter set for the Krasavietz class2.

| Par. | Value | Par. | Value |
|---|---|---|---|
| $\Delta$ t | 0.001 | $\tau$ | 0.008 |
| $afn$ | 3.6650390625 | $afp$ | -13.3037109375 |
| $bfn$ | -1.4853515625 | $cfn$ | 4.1650390625 |
| $agn$ | 1.6240234375 | $agp$ | 14.0205078125 |
| $bgn$ | -0.6923828125 | $cgn$ | 11.5205078125 |
| $ahn$ | -0.603515625 | $ahp$ | 5.62890625 |
| $bhn$ | -4.0517578125 | $chn$ | 4.6513671875 |
| $I_{b0}$ | 2.236328125 | $k_I$ | 16.703125 |
| $\phi$ | 0.19921875 | $\epsilon_q$ | 0.01171875 |
| $r_g$ | -1.126953125 | $r_h$ | -1.6884765625 |
| $m_0$ | -1 | $m_1$ | 1 |

Table 3: Parameter set for the NP1227 class1.

| Par. | Value | Par. | Value |
|---|---|---|---|
| $\Delta$ t | 0.001 | $\tau$ | 0.008 |
| $afn$ | 1.126953125 | $afp$ | -10.0498046875 |
| $bfn$ | -1.189453125 | $cfn$ | -8.6083984375 |
| $agn$ | -1.2724609375 | $agp$ | 13.484375 |
| $bgn$ | -6.3671875 | $cgn$ | 2.560546875 |
| $ahn$ | -0.78125 | $ahp$ | 11.970703125 |
| $bhn$ | -3.76953125 | $chn$ | 4.0849609375 |
| $I_{b0}$ | -8.587890625 | $k_I$ | 32.14453125 |
| $\phi$ | 1.3974609375 | $\epsilon_q$ | 0.0107421875 |
| $r_g$ | -1.7109375 | $r_h$ | -1.6904296875 |
| $m_0$ | -1 | $m_1$ | 1 |

[2] H. Kazama and R. I. Wilson, "Homeostatic matching and nonlinear amplification at identified central synapses," Neuron, vol. 58, no. 3, pp. 401–413, 2008. [Online]. Available: https://www.sciencedirect.com/science/article/pii/S0896627308001840

[3] N. K. Tanaka, H. Tanimoto, and K. Ito, "Neuronal assemblies of the drosophila mushroom body," Journal of Comparative Neurology, vol. 508, no. 5, pp. 711–755, 2008.

[4] O. Barnstedt, D. Owald, J. Felsenberg, R. Brain, J.-P. Moszynski, C. Talbot, P. N. Perrat, and S. Waddell, "Memory-relevant mushroom body output synapses are cholinergic," Neuron, vol. 89, no. 6, pp. 1237–1247, 2016.

[5] Y. Shang, A. Claridge-Chang, L. Sjulson, M. Pypaert, and G. Miesenbock, "Excitatory local circuits and their implications for olfactory processing in the fly antennal lobe," Cell, vol. 128, pp. 601–12, 03 2007.

[6] A. Das, A. Chiang, S. Davla, R. Priya, H. Reichert, K. Vijayraghavan, and V. Rodrigues, "Identification and analysis of a glutamatergic local interneuron lineage in the adult drosophila olfactory system," Neural systems and circuits, vol. 1, p. 4, 01 2011.

[7] S. Olsen, V. Bhandawat, and R. Wilson, "Excitatory interactions between olfactory processing channels in the drosophila antennal lobe," Neuron, vol. 54, pp. 89–103, 05 2007.

Table 4: Parameter set for the NP2426 class1.

| Par. | Value | Par. | Value |
|---|---|---|---|
| $\Delta$ t | 0.001 | $\tau$ | 0.008 |
| $afn$ | 5.279296875 | $afp$ | -15.05859375 |
| $bfn$ | -0.94921875 | $cfn$ | -1.0498046875 |
| $agn$ | -14.443359375 | $agp$ | 6.1220703125 |
| $bgn$ | -4.09375 | $cgn$ | 15.2958984375 |
| $ahn$ | -4.63671875 | $ahp$ | -0.97265625 |
| $bhn$ | -3.5986328125 | $chn$ | 4.21484375 |
| $I_{b0}$ | -13.65234375 | $k_I$ | 11.9833984375 |
| $\phi$ | 0.5576171875 | $\epsilon_q$ | 0.046875 |
| $r_g$ | -3.203125 | $r_h$ | -4.390625 |
| $m_0$ | -1 | $m_1$ | 1 |

Table 5: Parameter set for the PN.

| Par. | Value | Par. | Value |
|---|---|---|---|
| $\Delta$ t | 0.001 | $\tau$ | 0.00390625 |
| $afn$ | 0.25 | $afp$ | -0.25 |
| $bfn$ | -3.0 | $cfn$ | 0.0 |
| $agn$ | 0.125 | $agp$ | 1.0 |
| $bgn$ | -2.0 | $cgn$ | -4.0 |
| $\phi$ | 3 | $\theta$ | 0.0078125 |
| $r_g$ | -0.5 | $k_I$ | 1 |
| $k_0$ | -0.5 | $k_1$ | -4 |
| $k_r$ | 12 | $I_{b0}$ | -4 |
| $I_{b1}$ | -16 | $m_0$ | -1 |
| $m_1$ | 1 | $F_t$ | 1 |
| $\kappa$ | 0.00390625 | $m_0$ | -1 |
| $m_1$ | 4 | | |

[8] F. Python and R. F. Stocker, "Immunoreactivity against choline acetyltransferase, $\gamma$-aminobutyric acid, histamine, octopamine, and serotonin in the larval chemosensory system of dosophila melanogaster," Journal of Comparative Neurology, vol. 453, no. 2, pp. 157–167, 2002.

[9] R. I. Wilson and G. Laurent, "Role of gabaergic inhibition in shaping odor-evoked spatiotemporal patterns in the drosophila antennal lobe," Journal of Neuroscience, vol. 25, no. 40, pp. 9069–9079, 2005.

[10] S. Olsen and R. Wilson, "Lateral presynaptic inhibition mediates gain control in an olfactory circuit," Nature, vol. 452, pp. 956–60, 05 2008.

[11] C. M. Root, K. Masuyama, D. S. Green, L. E. Enell, D. R. Nassel, C.-H. Lee, and J. W. Wang, "A presynaptic gain control mechanism fine-tunes olfactory behavior," NEURON, vol. 59, no. 2, pp. 311–321, 7 2008.

[12] Y. Aso, D. Hattori, Y. Yu, R. M. Johnston, N. A. Iyer, T.-T. Ngo, H. Dionne, L. Abbott, R. Axel, H. Tanimoto, and G. M. Rubin, "The neuronal architecture of the mushroom body provides a logic for associative learning," eLife, vol. 3, 2014.

Table 6: Parameter set for the KC.

| Par. | Value | Par. | Value |
|---|---|---|---|
| $\Delta$ t | 0.001 | $\tau$ | 0.00390625 |
| $afn$ | 0.25 | $afp$ | -0.25 |
| $bfn$ | -3.0 | $cfn$ | 0.0 |
| $agn$ | -0.25 | $agp$ | 1.0 |
| $bgn$ | -4.0 | $cgn$ | -4.0 |
| $\phi$ | 2 | $\theta$ | 0.0625 |
| $r_g$ | -0.5 | $k_I$ | 1 |
| $k_0$ | -0.5 | $k_1$ | -4 |
| $k_r$ | 12 | $I_{b0}$ | -4.2 |
| $I_{b1}$ | -19.52 | $m_0$ | -1 |
| $m_1$ | 1 | | |

Table 7: Parameter set for the MBON.

| Par. | Value | Par. | Value |
|---|---|---|---|
| $\Delta$ t | 0.001 | $\tau$ | 0.00390625 |
| $afn$ | 1.0 | $afp$ | -0.25 |
| $bfn$ | -1.0 | $cfn$ | 0.0 |
| $agn$ | -0.25 | $agp$ | 1.0 |
| $bgn$ | -3.0 | $cgn$ | -4.0 |
| $\phi$ | 0.5 | $\theta$ | 0.03125 |
| $r_g$ | -0.5 | $k_I$ | 4 |
| $k_0$ | -0.5 | $k_1$ | -1 |
| $k_r$ | 4 | $I_{b0}$ | -4.8 |
| $I_{b1}$ | -2.3 | $m_0$ | -1 |
| $m_1$ | 2 | | |

Table 8: Parameter set for the APL.

| Par. | Value | Par. | Value |
|---|---|---|---|
| $\Delta$ t | 0.001 | $\tau$ | 0.00390625 |
| $afn$ | 0.125 | $afp$ | -0.125 |
| $bfn$ | -0.0 | $cfn$ | 0.0 |
| $agn$ | 0 | $agp$ | 1.0 |
| $bgn$ | -0.0 | $cgn$ | -4.0 |
| $I_{b0}$ | -5 | $k_I$ | 1 |
| $\phi$ | 0.125 | $\epsilon_q$ | 0.0078125 |
| $r_g$ | -0.0 | $r_h$ | 0 |
| $m_0$ | -1 | $m_1$ | 1 |

Table 9: Parameter set for the SMP354.

| Par. | Value | Par. | Value |
|---|---|---|---|
| $\Delta$ t | 0.001 | $\tau$ | 0.00390625 |
| $afn$ | 0.25 | $afp$ | -0.25 |
| $bfn$ | -3.0 | $cfn$ | 0.0 |
| $agn$ | -0.25 | $agp$ | 1.0 |
| $bgn$ | -4.0 | $cgn$ | -4.0 |
| $ahn$ | 0 | $ahp$ | 0 |
| $bhn$ | 0 | $chn$ | 0 |
| $I_{b0}$ | -4.2 | $k_I$ | 1 |
| $\phi$ | 4 | $\epsilon_q$ | 0 |
| $r_g$ | -0.5 | $r_h$ | 0 |
| $m_0$ | -1 | $m_1$ | 1 |

Table 10: Values of scaling parameters $p_{x\_y}$.

| $x$ | $y$ | Value | $x$ | $y$ | Value |
|---|---|---|---|---|---|
| ORN | LN | 0.006591796875 | ORN | PN | 0.125 |
| LN | LN | 0.00390625 | LN | PN | 0.004150390625 |
| PN | LN | 1.0 | PN | PN | 0.125 |
| PN | KC | 1.03125 | PN | APL | 0.078125 |
| KC | MBON-$\alpha$3 | 0.3125 | KC | MBON-$\alpha$1 | 0.5625 |
| KC | APL | 0.078125 | APL | MBON-$\alpha$3 | 1 |
| APL | MBON-$\alpha$1 | 1 | MBON-$\alpha$3 | SMP354 | 0.28125 |
| MBON-$\alpha$1 | SMP354 | 0.28125 | | | |
